# Lymphatic CD49a is a driver of meningeal immune aging and cognitive decline

**DOI:** 10.64898/2026.08.04.742777

**Authors:** Natalie M. Frederick, Rachel Tinkey, Gabriel A. Tavares, Caitlyn N. Dahnke, Henry Busch, Nikhita Arun, Logan Chung, Andrew D. Buxbaum, Antoine Louveau

## Abstract

Aging is associated with progressive accumulation and dysregulation of dural immune cells, coinciding with impaired CSF drainage and lymphatic function. Prior work has shown that improving lymphatic function in aged mice is sufficient to ameliorate age-associated cognitive decline, and that local immune cells can directly regulate lymphatic draining function. Yet, the endothelial-intrinsic mechanisms driving lymphatic dysfunction remain unclear. Here we found that the integrin CD49a is upregulated in aged lymphatic endothelial cells and regulates CCL21 release. Accordingly, genetic deletion of CD49a in lymphatic endothelial cells broadly reverses age-associated immune dysfunction across dural myeloid, lymphoid and dendritic cell compartments, limits glial aging, and mitigates cognitive and social behavioral deficits, thereby revealing a targetable endothelial-intrinsic mechanism of lymphatic aging.

## Introduction

Aging is accompanied by a progressive, low-grade inflammation state within the central nervous system (CNS) that contributes to cognitive decline and increases susceptibility to neurodegenerative disease (Franceschi and Campisi, 2014; Barrientos et al., 2015; Deczkowska et al., 2018; Baruch et al., 2014). While much of this work has focused on parenchymal glial cells (Deczkowska et al., 2018; Norden and Godbout, 2013; Clarke et al., 2018; Labarta-Bajo and Allen, 2025; Segel et al., 2019; Dulken et al., 2019; Kaya et al., 2022), it is now clear that the meninges, and particularly the dura mater, harbor a diverse and dynamic immune compartment that both senses and shapes CNS immune status across the lifespan (Rustenhoven and Kipnis, 2022; Alves de Lima et al., 2020). With age, the dural immune landscape becomes progressively skewed toward chronically activated, exhausted and pro-inflammatory phenotypes across myeloid, lymphoid and dendritic cell populations, mirroring changes described in peripheral immunosenescence (Nikolich-Žugich, 2018; Krishnarajah et al., 2022; Mrdjen et al., 2018; Dulken et al., 2019). However, the endothelial and stromal mechanisms that permit, or actively drive, this age-associated remodeling remain poorly defined.

The dural lymphatic vasculature has become increasingly appreciated as a central regulator of CNS fluid and immune homeostasis (Louveau et al., 2018; Da Mesquita et al., 2018a; Zhao et al., 2025). These vessels drain cerebrospinal fluid (CSF), interstitial solutes and immune cells from the meninges and brain parenchyma to the cervical lymph nodes (CLN), providing a direct conduit linking the CNS to peripheral immune surveillance (Louveau et al., 2015, 2018; Aspelund et al., 2015). Dural lymphatic function declines with age, characterized by reduced vessel width and impaired macromolecule clearance (Da Mesquita et al., 2018b; Zhou et al., 2020). This decline coincides with an age-related loss of CCR7-dependent immune cell egress through the dural lymphatics, which independently contributes to glymphatic dysfunction and cognitive decline (Da Mesquita et al., 2021a). Restoration of lymphatic function in aged mice through Vegfc-induced lymphangiogenesis ameliorates age-associated cognitive decline, establishing dural lymphatic dysfunction as a causal, and therapeutically tractable, contributor to brain aging (Da Mesquita et al., 2018b, 2021b). Conversely, local immune cells can directly regulate lymphatic endothelial cells (LECs) function, as aging increases meningeal T cell numbers and IFNγ production. Mechanistically, IFNγ signaling on LECs disrupts VE-cadherin junctional integrity to impair CSF drainage (Rustenhoven et al., 2023), indicating bi-directional crosstalk between the dural immune and lymphatic compartments.

Integrins are heterodimeric transmembrane receptors that mediate cell-extracellular matrix (ECM) interactions and have emerged as key regulators of LECs identity and function, including valve formation and vessel maturation (Bazigou et al., 2009; Chen et al., 2014; Hitpass Romero et al., 2025a; Hong et al., 2004). Among the integrin family, CD49a (integrin α1, Itga1) forms the collagen-binding integrin α1β1 through heterodimerization with the β1 subunit (Itgb1), and has well-established roles in regulating immune cell phenotype, notably the establishment and maintenance of tissue-resident memory T cells (Ray et al., 2004; Bromley et al., 2020). In addition, CD49a is also broadly expressed by cells that contact vascular and perivascular basement membranes, including endothelial cells (Pang et al., 2023; Hong et al., 2004), but its expression and function on LECs in the dura has not been previously examined.

Here, we identify CD49a as an age-upregulated integrin in dural LECs and investigate its role in aging. Using lymphatic-specific deletion, we show that the loss of CD49a does not alter lymphatic vessel morphology or bulk fluid drainage but instead restores immune cell trafficking by promoting the secretory release of the chemokine CCL21. This endothelial-intrinsic change is sufficient to broadly reprogram the aged immune compartment of the dura, attenuate glial aging in the underlying cortex, and improve social and cognitive behavior in aged female mice. Together, these findings identify an intrinsic switch in LECs as a driver of CNS immune aging and establish CD49a as a candidate for restoring dural lymphatic and immune function in the aged brain.

## Results

### CD49a regulates immune cells drainage in aged LECs

To investigate how the dural LECs change in aging, we performed pathway analysis of a previously published bulk RNA sequencing dataset (Da Mesquita et al., 2018b). Based on the differentially expressed genes (DEGs), seven of the top 30 enriched pathways are related to the ECM, fibrosis and integrin signaling (Figure 1A), suggesting that aging remodels the interactions between dural LECs and their microenvironment. Integrins are the major class of cell surface receptors mediating cell-ECM interactions, and changes in ECM composition have been shown to alter dural lymphatic function (Hitpass Romero et al., 2025a). Analysis of an independent RNA-seq dataset (Louveau et al., 2018) shows that dural LECs express high levels of *Itga9*, a lymphatic associated integrin important for valves formation and function (Bazigou et al., 2009), as well as *Itga1* (CD49a) and *Itga11* (Supplementary Figure 1A). Comparing expression levels of all alpha- and beta-subunit isoforms revealed significant upregulation of *Itga1* and *Itgb1* selectively in aged dural LECs (Figure 1B). To validate these transcriptomic findings, we examined CD49a protein expression and found that, consistent with the RNA-seq data (Supplementary Figure 1A), dural and diaphragm LECs express higher level of CD49a than skin LECs (Figure 1C and D), suggesting a degree of tissue-specific enrichment of CD49a. Measurements of CD49a expression by dural LECs confirmed a significant upregulation of CD49a in aged LECs compared to younger mice (Figure 1E and F). Collectively, these results demonstrate that CD49a expression is enriched in dural LECs and exhibits age-associated upregulation.

**Figure 1:**
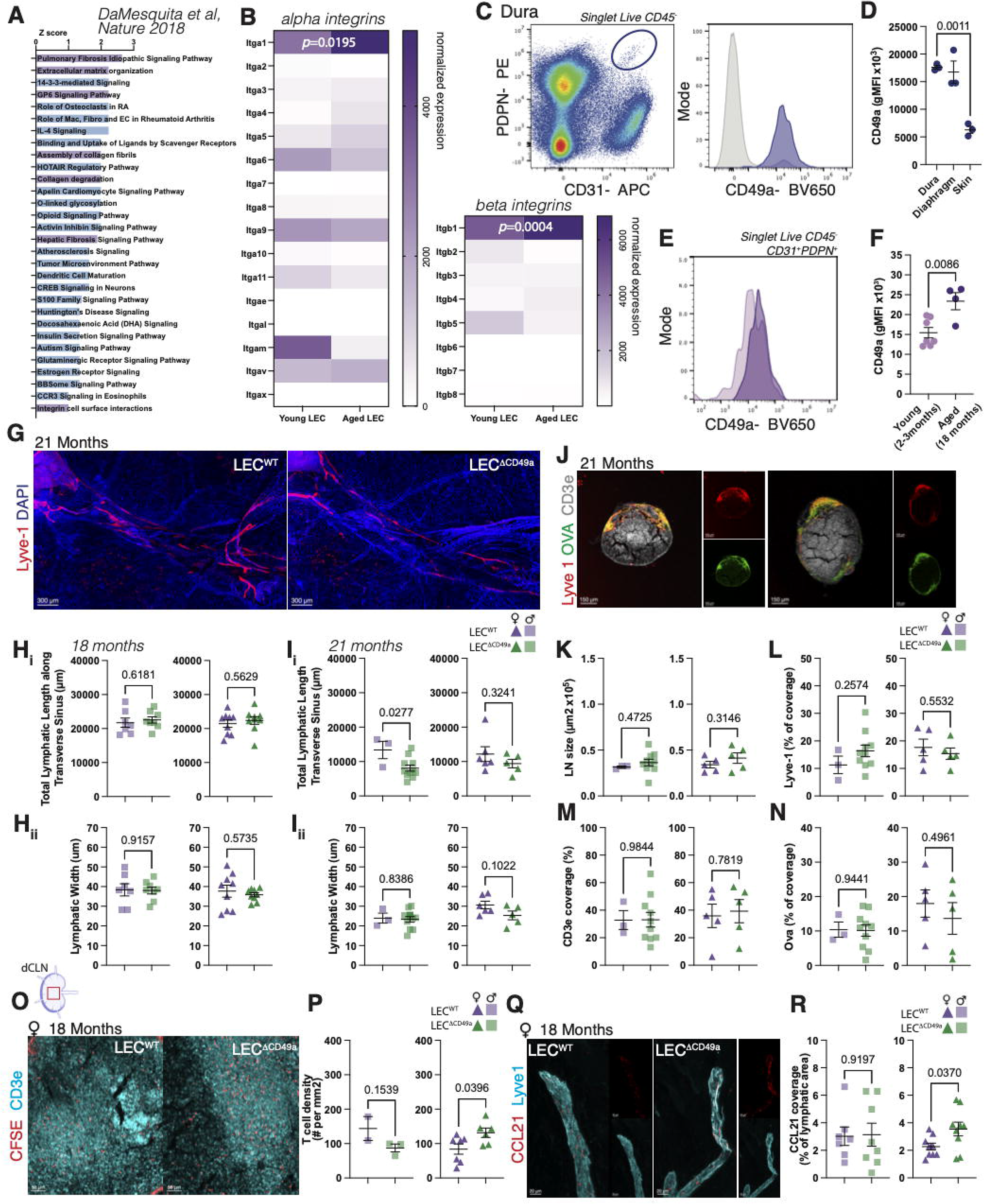
CD49a is upregulated in aging and loss of CD49a in lymphatic endothelial cells improves Ccl21 expression and immune cell drainage in aged mice.. A. Ranked significant pathway analysis comparing dural LECs from young and aged mice. B. Heatmap comparison of normalized expression of alpha and beta chain of integrin subunit in young and aged dural LECs. N=3 biological replicate per group. Two-way ANOVA with Sidak’s multiple comparisons test. C. Representative gating of lymphatic endothelial cells from the dura (Cd45-Cd31+Pdpn+) and histogram of CD49a expression in dural LECs (blue line) and double negative (Cd45-Cd31-Pdpn-) dural cells. D. Quantification of the geometric mean fluorescence intensity of CD49a expression in LECs from the dura, diaphragm and skin. Mean ± s.e.m. N=3 mice per group. One way ANOVA with Dunnett’s multiple comparisons test. E. Representative histogram of CD49a expression in dural LECs from young and aged mice. F. Quantification of the geometric mean fluorescence intensity of CD49a expression in LECs from the dura of young and aged mice. Mean ± s.e.m. Unpaired t test. G. Representative images of the dural lymphatic vessels (Lyve-1;red) along the transverse sinus of aged (21 months old) mice. H. Quantification of the total lymphatic length (i) and lymphatic width (ii) along the transverse sinus of 18 months old male (square) and female (triangle) mice. Mean ± s.e.m. Unpaired t test. I. Quantification of the total lymphatic length (i) and lymphatic width (ii) along the transverse sinus of 21 months old male (square) and female (triangle) mice. Mean ± s.e.m. Unpaired t test. J. Representative images of CSF-injected OVA (green) in the dCLNs of 21 months old mice. Lymph nodes are characterized by the presence of a capsule (Lyve-1; red) and T cells zone (Cd3e; gray). K. Quantification of the size of the lymph node section. Mean ± s.e.m. Unpaired t test. L. Quantification of the percentage of Lyve1 coverage. Mean ± s.e.m. Unpaired t test. M. Quantification of the CD3e coverage. Mean ± s.e.m. Unpaired t test. N. Quantification of the OVA coverage. Mean ± s.e.m. Unpaired t test. O. Representative images of CSF injected CFSE labelled naïve CD4 T cells in the T cells zone of the dCLNs of 18 months old mice. P. Quantification of the density of CFSE+ T cells in the dCLNs. Mean ± s.e.m. Unpaired t test. Q. Representative images of the chemokine Ccl21 (red) in the dural lymphatic vessels (Lyve-1; cyan) of 18 months old mice. R. Quantification of the Ccl21 coverage of the dural lymphatic vessels. Mean ± s.e.m. Unpaired t test.

To examine the role of CD49a in LECs, we crossed CD49a floxed mice with Prox1creERT2 mice (Supplementary Figure 1B). Tamoxifen injection at 80mg/kg for 4 days at 6 weeks led to sustained and specific deletion of CD49a from LECs, sparing blood endothelial cells (Supplementary Figure 1 C and D). In adult mice, *Prox1* is also expressed in neurons of the dendate gyrus (Lavado and Oliver, 2007; Galeeva et al., 2007); however, CD49a expression was detected in neurons of the hippocampus (Supplementary Figure 1E). Analysis of dural lymphatic morphology at 3, 18 and 21 months of age revealed no significant effects of CD49a deletion on lymphatic length or width regardless of age or sex (Figure 1G to I, Supplementary Figure 1F and G). Similarly, measurement of cerebrospinal fluid (CSF) injected ovalbumin as a proxy of dural lymphatic fluid drainage capacity (Louveau et al., 2015) demonstrated no change with CD49a deletion at any age (Figure 1J to N; Supplementary Figure 1H to L).

Lymphatic vessels don’t only drain fluid, they also facilitate immune cell trafficking and recirculation (Randolph et al., 2017), a process that declines with aging (Kataru et al., 2022; Zolla et al., 2015). Therefore, we next assessed whether CD49a deletion affected the immune cell drainage capacity of the dural lymphatics. Naïve CD4 T cells isolated from the spleen, which express high levels of CCR7 and serve as a model for CCR7-dependent immune cell trafficking (Louveau et al., 2018) were injected into the CSF. After 12h, their accumulation in the deep cervical lymph nodes (dCLNs) was quantified, as previously described (Louveau et al., 2018). In young mice, the deletion of CD49a from LECs did not alter the drainage of immune cells into the dCLNs (Supplementary Figure 1M and N). However, at 18 months, we found an increased density of injected immune cells in female mice lacking CD49a expression in LECs (Figure 1 O and P). There were no observable differences in male mice (Figure 1 O and P). Given that CCL21 expression and release by LECs is one of the major drivers of immune cell drainage (Louveau et al., 2018; Förster et al., 2008; Bromley et al., 2005), we next measured CCL21 expression by dural LECs. In accordance with the immune drainage data, we found that female mice showed increased levels of CCL21 at 18 months of age (Figure 1Q and R), with no difference in aged male mice (Figure 1Q and R), or in young mice (Supplementary Figure 1O and P). Overall, our data demonstrates that CD49a is upregulated in aged LECs and that LEC-specific loss of CD49a enhances CCL21 expression and promotes associated immune cell trafficking in an age-and sex-dependent manner.

### CD49a partially regulates the transcriptome of aged dural LECs

To better understand how CD49a regulates LEC function, we sorted the CD31+ cells from the dura of young and aged mice with conditional knockout (KO) of CD49a in LECs and littermate controls (Figure 2A). Cell clustering of transcriptomic profiles following quality control identified four main endothelial populations (Figure 2B) corresponding to 3 blood endothelial clusters and one LECs cluster. LECs were identified based on the expression of canonical markers such as *Prox1*, *Lyve1* and *Flt4* (Figure 2C). Consistent with findings in Figure 1, CD49a transcript expression in dural LECS increased in aged LECs and was decreased in the conditional KO, with limited changes observed in blood endothelial cells (Figure 2D). Comparison of the transcriptomic profile of dural LECs from young mice conditional CD49a KO mice and littermate controls revealed no significant transcriptional changes, while in aged mice, we observed 117 DEGs, comprising 91 upregulated and 26 downregulated genes (Supplementary Table 1). These results suggest that CD49a has a stronger impact on the transcriptomic profile of LECs in aged mice compared to that of young mice. Furthermore, functional categorization of DEGs highlighted that loss of CD49a altered pathways associated with LEC homeostasis, including mitochondrial function, response to oxidative stress, proteostasis, transcription, translation and RNA processing (Figure 2E), all of which are hallmarks of aging cells (López-Otín et al., 2013, 2023). Numerous genes dysregulated by the loss of CD49a were also associated with vesicular transport and cytoskeletal organization (Figure 2E).

**Figure 2:**
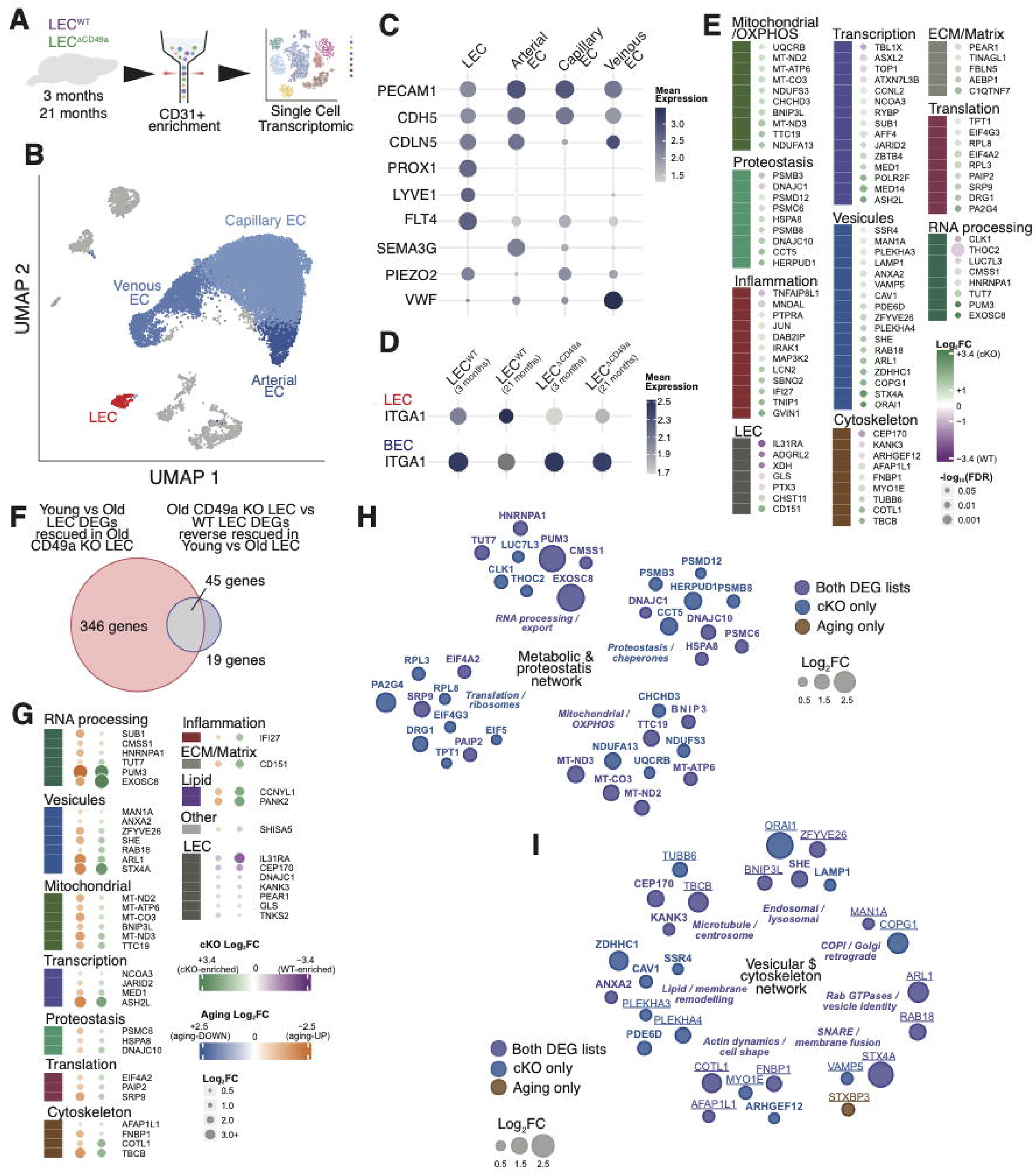
CD49a regulates the transcriptomic profile of aged dural LECs. A. Schematic of the experiment. Cd31+ cell fraction from the dura of young and aged mice were FACS sorted and sequenced using the 10x genomic platform. B. UMAP projection of single-cell RNA-seq data colored by cell type. Grey clusters are unassigned non-endothelial cells. C. Bubble plot of mean expression of vascular markers by the different clusters. D. Bubble plot of *Itga1* (CD49a) mean expression in lymphatic endothelial cells (LECs) and blood endothelial cells (BEC) across sample groups. E. Heatmap-style list of differentially expressed genes between aged LEC^ΔCD49a^ and LEC^WT^ grouped by functional category. Color indicate Log2fold-change and dot size indicates statistical significance (-log10FDR). F. Venn diagram comparing two DEG lists: genes differentially expressed between young and old LECs that are rescued in aged LEC^ΔCD49a^, and genes differentially expressed between old LECS^ΔCD49a^ and LEC^WT^ whose direction is reverse related to the young-vs-old aging signature. 45 genes overlap between the two lists, representing aging-associated changes that are rescued by the loss of CD49a. G. Dot plot of the 45 overlapping genes from (F) grouped by functional category, displaying both cKO Log2FC (green-purple scale) and aging Log2FC (blue-orange scale) side by side, with dot size representing magnitude of fold change. H. Gene interaction network of the “Metabolic and proteostasis” module. Node color denotes whether gene appears in both DEG lists (purple), the cKO-only list (blue) or the aging-only list (brown); node size reflects Log2FC magnitude. I. Gene interaction network of the “Vesicular and cytoskeleton network” module. Node color denotes whether gene appears in both DEG lists (purple), the cKO-only list (blue) or the aging-only list (brown); node size reflects Log2FC magnitude.

To determine whether genotype-associated CD49a rescues the transcriptional signature of physiological aging, we generated an “aging” DEGs signature by comparing the transcriptomic profile of LECs from aged mice to young mice, and the genotype associated signature (Figure 2E). For each significant DEG identified in either list, we extracted the corresponding fold change (irrespective of statistical significance) from the other comparison, allowing us to assess whether genotype-driven changes trend in the same or opposite direction as physiological aging. The intersection of the two analyses revealed that 45 DEGs showed concordant fold changes and were defined as a core set of genes whose regulation by CD49a loss opposes the transcriptional trajectory of normal aging (Figure 2F), suggesting that CD49a deletion partially rescues the transcriptomic profile of aged LECs. Organizing these genes by cellular function highlighted similar pathways to those of the DEGs from the conditional CD49a KO LECs (Figure 2E), including RNA processing, mitochondrial function, transcription, translation, and proteostasis, as well as vesicular transport and cytoskeletal organization (Figure 2G). Moreover, this analysis identified alterations in two major biological networks following CD49a loss, including metabolic and proteostasis pathways as well as vesicular trafficking and cytoskeletal architecture (Figure 2H and I). Interestingly, changes in the metabolic and proteostasis networks highlighted a general upregulation of immunoproteasome-associated genes, such as *Psmd12*, *Psmb8*, and *PsmcC* suggesting a heightened capacity of aged LECs to degrade oxidatively damaged and misfolded proteins upon loss of CD49a (Seifert et al., 2010a; Vilchez et al., 2012). Genes associated with RNA metabolism, processing and transport were also upregulated following CD49a deletion, consistent with changes in RNA homeostasis in aged LECs. In addition, increased expression of mitochondrial-associated genes such as *Ttc19* (to prevent oxidative damage (Ghezzi et al., 2011; Bottani et al., 2017)), *Chchd3* (inner mitochondrial membrane regulator supporting ATP generating machinery (Darshi et al., 2011)) and *Bnip3l* (mitophagy promoter (Schweers et al., 2007; Novak et al., 2010)), suggest improved mitochondrial function. CCL21 release depends on a multi-step, tightly regulated trafficking pathway rather than passive diffusion, as it is stored intracellularly within Golgi-derived vesicles and is primarily triggered locally by a calcium signal generated upon contact with a transmigrating cell (Vaahtomeri et al., 2017; Russo et al., 2016). As a result, we hypothesized that the vesicular trafficking and cytoskeletal changes observed in our DEG analysis could impair the CCL21 secretory capacity of LECs in the conditional KO, leading to increased intracellular retention of CCL21 (Figure 1Q and R). Additionally, because we observed sex-specific alterations in lymphatic drainage *in vivo*, we assessed whether the transcriptomic changes exhibited similar sex biases. We bioinformatically partitioned cells based on their expression of the Y-chromosome genes *Uty* and *Ddx3y* (Chen et al., 2016). All but two genes (*Eif4g3* and *Man1a*) showed similar directionality in fold change between male and female cells (Supplementary Figure 2A), suggesting that loss of CD49a induces a similar transcriptomic response in LECs regardless of sex. We also confirmed that the vesicular and cytoskeletal signature is present in the bulk RNA-seq, comparing wildtype aged and young LECs (Supplementary Figure 2B), demonstrating that changes in vesicular transport are a feature of aging meningeal LECs.

### CD49a activation in vitro directly regulates ATP-dependent transport of CCL21

To better understand how CD49a regulates LECs metabolic function and vesicular trafficking, we utilized an *in vitro* model of primary, commercially available, LECs isolated from young and aged mice. qPCR analysis confirmed that both young and aged cultures maintained a comparable LEC transcriptional profile, with no detectable expression of non-LEC markers (Supplementary Figure 3A). Following validation of LEC identity, we assessed CD49a expression and found that, consistent with our *in vivo* findings, aged cells expressed significantly higher levels of CD49a than young cells by both flow cytometry (Figure 3A and B) and western blot analysis (Figure 3C and D). With this understanding, we next attempted to recapitulate a microenvironment of chronic CD49a activation to assess the functional output. The Ha31/8 clone of anti-CD49a antibody has traditionally been used as a functional inhibitor of CD49a, based on its ability to promote detachment of cells cultured on collagen IV substrates (Mendrick et al., 1995). However, we hypothesized that antibody engagement of CD49a may also directly induce receptor activation in treated cells. To test this, LECs were cultured on a gelatin solution composed of denatured collagen I fibers that are not a main ligand for CD49a to limit integrin-based adhesion (Davis, 1992; Knight et al., 2000). Young and aged LECs were subsequently stimulated with Ha31/8 to evaluate CD49a activation. We found that both focal adhesion kinase (FAK) and SRC family proteins were phosphorylated as early as 30 minutes after stimulation in young LECs compared to IgG-treated controls (Supplementary Figure 3B and C), demonstrating activation of integrin signaling. Interestingly, aged LECs, however, appeared unresponsive to anti-CD49a treatment with no detectable long-term changes in phosphorylation of FAK or SRC (Supplementary Figure 3B and C). Overall, these data demonstrate that the Ha31/8 clone of anti-CD49a indeed activates CD49a in young LECs.

**Figure 3:**
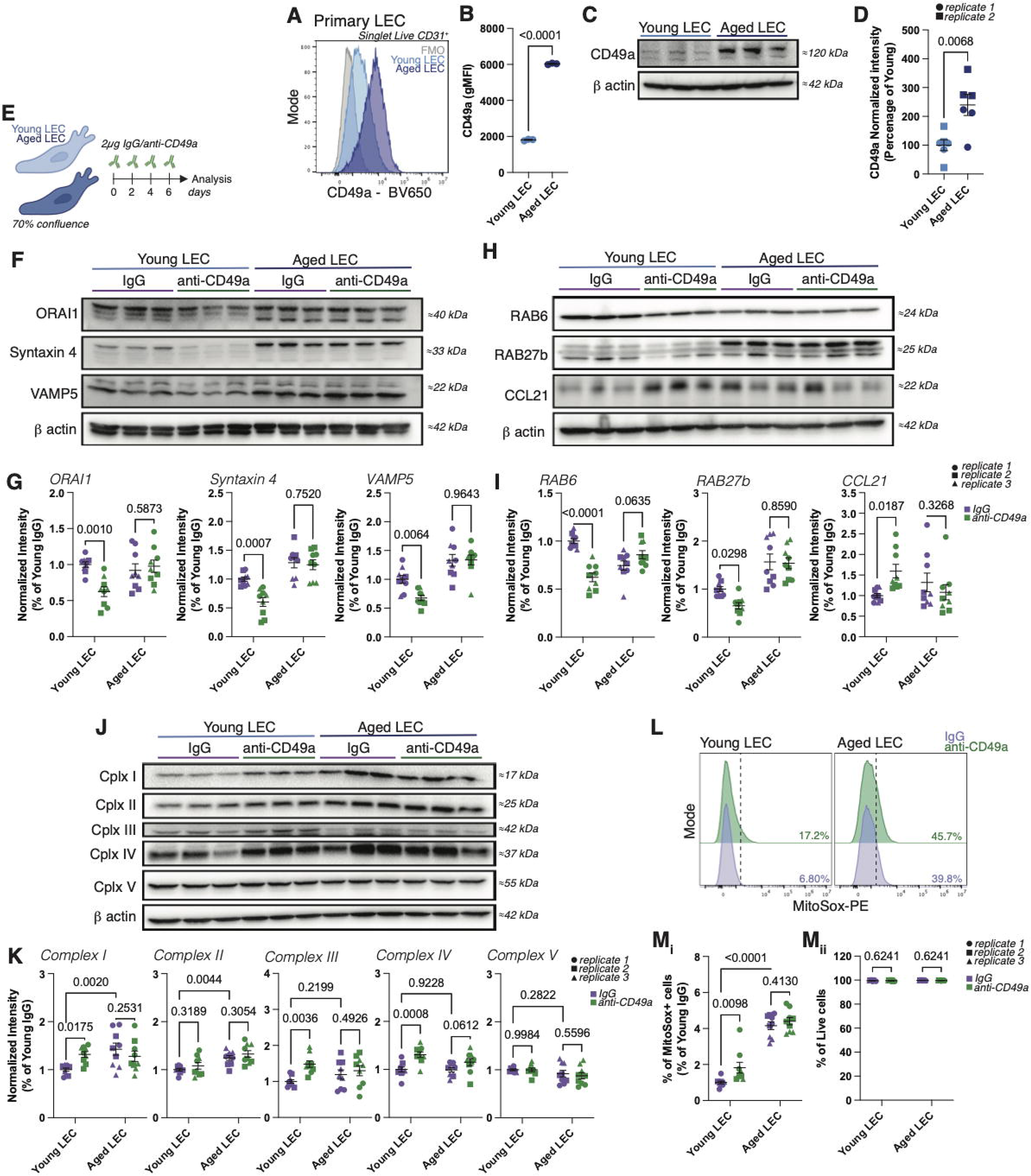
CD49a directly regulate Ccl21 trafficking and mitochondrial function in LECs. A. Representative histogram of CD49a expression in young and aged LECs *in vitro*. B. Quantification of the mean fluorescence intensity of CD49a expression in young and aged LECs in vitro. Mean ± s.e.m. Unpaired t test. C. Representative western blot panels of CD49a and beta actin expression in young and aged LECs *in vitro*. D. Quantification of the normalized CD49a expression in young and aged LECs *in vitro*. Samples from two replicate experiments are shown and coded by shape. Mean ± s.e.m. Unpaired t test. E. Schematic of chronic CD49a activation *in vitro*. Young and aged LECs at 70% confluence were treated every other day with 2µg of IgG or anti-CD49a in a 96 well plate. Cells were harvested at day 7 after treatment initiation. F. Representative western blot panels for Orai1, Syntaxin 4, Vamp5 and beta Actin in young and aged LECS treated with IgG or anti-CD49a. G. Quantification of the normalized expression of Orai1, Syntaxin 4 and Vamp5 expressed as a percentage of the young IgG condition. Samples from 3 independent replicate experiment are shown and coded by shape. Mean ± s.e.m. Two-way ANOVA with Uncorrected Fisher’s LSD. H. Representative western blot panels for Rab6, Rab27b, Ccl21 and beta Actin in young and aged LECs treated with IgG or anti-CD49a. I. Quantification of the normalized expression of Rab6, Rab27b and Ccl21 expressed as a percentage of the young IgG condition. Samples from 3 independent replicate experiments are shown and coded by shape. Mean ± s.e.m. Two-way ANOVA with Uncorrected Fisher’s LSD. J. Representative western blot panels mitochondrial respiratory chain complexes I to V and beta actin in young and aged LECs treated with IgG or anti-CD49a. K. Quantification of the normalized expression of mitochondrial respiratory chain complexes expressed as a percentage of the young IgG condition. Samples from 3 independent replicate experiments are shown and coded by shape. Mean ± s.e.m. Two-way ANOVA with Uncorrected Fisher’s LSD. L. Representative histogram of MitoSox staining in young and aged LECs treated with IgG or anti-CD49a. M. Quantification of the percentage of Mitosox positive cells expressed as a percentage of the young IgG condition (i) and percentage of live cells (ii) in young and aged LECs treated with IgG or anti-CD49a. Samples from 3 independent replicate experiments are shown and coded by shape. Mean ± s.e.m. Two-way ANOVA with Uncorrected Fisher’s LSD.

To further investigate how chronic activation of CD49a affects LEC function, cultured LECs were stimulated every other day with Ha31/8 (or IgG control) for a duration of 7 days. We found that chronic CD49a activation led to the coordinated reduction of Orai1, Syntaxin 4 (Stx4) and Vamp5 (Figure 3F and G), 3 targets found to have rescued expression with LEC-specific deletion of CD49a in aged mice (Figure 2E). These proteins are components of the calcium-dependent, SNARE-mediated exocytosis machinery (Jahn and Fasshauer, 2012) broadly implicated in regulating vesicular secretion in various cell types (Vig et al., 2008; Olson et al., 1997; Zeng et al., 1998). Additionally, we found downregulation of two RAB GTPases previously identified as markers of CCL21 storage vesicles (Vaahtomeri et al., 2017), RAB6 (Golgi-derived vesicles) and RAB27b (dense-core secretory granules) (Figure 3H and I). This coordinated downregulation of vesicle-storage and fusion-machinery components was accompanied by an intracellular accumulation of CCL21 (Figure 3H and I), in the absence of transcriptomic changes (Supplementary Figure 3D), suggesting a block of its trafficking and/or release. Although CD49a loss increased intracellular CCL21 in aged mice *in vivo* (Figure 1Q and R), chronic CD49a activation increased, rather than decreased, intracellular CCL21 in cultured LECs. We propose that these apparently divergent results reflect the same underlying effect in vesicular trafficking capacity, manifesting differently depending on physiological demand for CCL21 release. *In vitro*, due to the absence of immune cells, CCL21 is secreted at a lower rate (Vaahtomeri et al., 2017; Kriehuber et al., 2001; Johnson and Jackson, 2010), and the impaired trafficking machinery caused by chronic CD49a activation leads to the accumulation of newly synthesized CCL21 that cannot be efficiently packaged or released. In contract, LEC *in vivo*, continuously maintain interstitial and haptotactic CCL21 gradients, necessitating ongoing replenishment of intracellular CCL21 pools (Sarris et al., 2012; Weber et al., 2013). Thus, impaired vesicular trafficking may limit CCL21 replenishment and vesicle biogenesis, allowing consumption to outpace production. This results in reduced intracellular CCL21 availability and is consistent with the loss of CCL21 expression observed in aged LECs compared to young LECs *in vivo*. Overall, these findings indicate that CD49a expression and activation in LECs disrupts the packaging and trafficking of CCL21, and that genetic loss of CD49a is sufficient to restore this expression and improved immune cell drainage in aged mice (Figure 1O and P). This establishes a framework in which CD49a-driven impairment of the CCL21 secretory pathway contributes to the age-associated decline in CNS lymphatic drainage.

In conjunction with CCL21 secretion, SNARE mediated vesicular release also requires adenosine triphosphate (ATP) (Cipriano et al., 2013; Jahn and Fasshauer, 2012). Notably, our RNA sequencing data suggested that the loss of CD49a affects mitochondrial gene expression (Figure 2H), providing a potential link between CD49a and the generation of ATP. To determine how CD49a affects mitochondrial function, we assessed whether there were differences in the expression of electron transport chain proteins. We found that chronic activation of CD49a in young LECs altered the expression profile of mitochondrial complexes with upregulation of complex I, III and IV and no change in complex II and V (Figure 3J and K). Measurement of total mitochondria using TOM20 and VDAC showed no differences following anti-CD49a treatment (Supplementary Figure 3E and F), demonstrating that the changes in complex proportions were not due to increased mitochondrial mass. These results were consistent with our transcriptomic analysis, which identified increased expression of *Uqcrb* and *Ttc19* in the conditional KO, genes associated with mitochondrial complex III (Ghezzi et al., 2011; Jung et al., 2010), as well as *Ndufs3* and *Ndufa13,* which are associated with complex I (Huang et al., 2004; D’Angelo et al., 2021). Functionally, changes in the expression of mitochondrial complexes could reflect different statuses of mitochondrial health and function (Guan et al., 2022; Ramírez-Camacho et al., 2020). Therefore, we measured the production of reactive oxygen species (ROS) upon anti-CD49a treatment. Chronic treatment of young LECs with anti-CD49a led to an increased percentage of MitoSox-positive LECs (Figure 3L and M), suggesting heightened production of ROS when CD49a is activated. Aged LECs showed a higher baseline level of ROS production (Figure 3L and M), correlating with increased expression of mitochondrial complex I, a major site of ROS production (Murphy, 2009; Cochemé and Murphy, 2008). Importantly, neither cell age nor CD49a activation altered the percentage of cell death, indicating that CD49a agonism does not affect LEC viability (Figure 3M). Together, these results demonstrate that CD49a activation promotes mitochondrial dysfunction and increased ROS production and is consistent with our transcriptomic findings. Conversely, loss of CD49a partially rescues the age-associated mitochondrial transcriptome in dural LECs, inferring a beneficial effect on CCL21 secretion via improved energy production. Collectively, our model suggests that CD49a may both directly (via regulation of vesicular trafficking components) and indirectly (via mitochondrial dysfunction) regulate the secretion of CCL21 by LECs.

Loss of CD49a in LECs improved the profile of the aged dural immune compartment.

To investigate how LEC-specific CD49a deletion and associated improvements in immune cell drainage affect the dural immune compartment of aged mice, we performed single-cell RNA sequencing. CD45+ cells from the dura of aged conditional CD49a KO and littermate controls were sorted and sequenced using the 10x genomic platform (Figure 4A). Unsupervised clustering identified all major dural immune lineages (Mrdjen et al., 2018; Van Hove et al., 2019; Korin et al., 2017), including neutrophils, myeloid cells (macrophages, monocytes and dendritic cells), B cells, lymphocyte lineages and innate lymphoid cells (Figure 4B), with cell identities assigned using canonical lineage markers (Supplementary Figure 4A). Comparison of conditional KO and WT mice demonstrated no changes in the presence of major immune cell populations (Figure 4C) but rather suggested a mild change in cell composition with increased neutrophils and B cells in the conditional KO compared to controls (Figure 4D). We therefore performed lineage-specific analysis to determine how CD49a loss alters individual immune cell populations. Sub-clustering of neutrophils identified a continuum spanning from promyelocyte-like and myelocyte-like progenitors through conventional mature and matrix-remodeling neutrophils to a transcriptionally distinct “inflammatory aged neutrophils state” characterized by the high expression of *Il1*β, and *Nlrp3* (Van Bruggen et al., 2023; Van Avondt et al., 2023; Sreejit et al., 2022) (Figure 4E and F). Proportion analysis showed a decrease in “inflammatory aged” neutrophils in the conditional CD49a KO, with a concomitant increase in the conventional and matrix-remodeling neutrophils populations (Figure 4G). Further, DEG analysis of “inflammatory aged” neutrophils identified decreased expression of immediate-early genes (*Fos*, *Junb*) and inflammatory-associated markers (*Il1b*, *Cles4d*, *Srgn*) in aged conditional KO mice, consistent with reduced activation of neutrophil state (Figure 4H, Supplementary Table 2). Similarly, sub-clustering of the T/NK/innate lymphoid compartment resolved distinct lymphoid populations, including ILC2s, γδ T cells, NK cells, regulatory T cells, exhausted/chronically activated CD4 T cells, tissue-resident memory CD8 T cells, exhausted/effector memory CD8 T cells, and IFN-stimulated T cells populations (Figure 4I and J). In addition, CD49a loss was also associated with a downregulation of mitochondrially encoded electron transport chain genes (*Mt-co3*, *Mt-nd1*, *Mt-nd2*, *Mt-nd3*, *Mt-atp6*, *Mt-cytb*) alongside increased expression of ribosomal and translation-associated transcripts (*Rpl* and *Rps* family genes, *Eif1*, *Uba52*) (Figure 4K to M, Supplementary Table 2), suggesting altered metabolic and translational profiles within these cells. Furthermore, several DEGs were indicative of a shift in T cell state. Notably, *Cblb* and *Ubash3b,* negative regulators of TCR signaling that are associated with T cell exhaustion (Kumar et al., 2021; Mikhailik et al., 2007), were downregulated in conditional KO mice (Figure 4L, Supplementary Table 2), suggesting reduced engagement of exhaustion-associated pathways. Likewise, upregulation of *Sh2d1a*, a critical mediator of NK/T cell cytotoxic effector function (Sharifi et al., 2004), together with *Psmb8,* which supports proteostasis under oxidative stress (Seifert et al., 2010b), was consistent with preservation of T cell function (Figure 4L, Supplementary Table 2). Additionally, we also observed altered expression of *Cd37* and *Ltb*, and while not directly associated with effector function, implies maintenance of T cell responses (Gartlan et al., 2013; Summers deLuca et al., 2011) (Figure 4L, Supplementary Table 2). Further analysis subclustered dural B cells into several distinct populations, including age-associated memory and proliferative B cells (Supplementary Figure 4B and C). Beyond a shared mitochondrial signature, age-associated memory B cells in the conditional CD49a KO mice showed reduced expression of *Zeb2*, the transcription factor required for age-associated B cell differentiation (Dai et al., 2024), along with reduced *Dock2* (Ushijima et al., 2018), suggesting that CD49a loss in LECs may attenuate the age-associated differentiation program (Supplementary Figure 4D, Supplementary Table 2). Additionally, although not among the highest-ranked DEGs by effect size, *Itgam* and *Cd80* were also significantly downregulated in the conditional CD49a KO. These changes are consistent with a shift away from the inflammatory age-associated B cell phenotype (Masle-Farquhar et al., 2022) (Supplementary Table 2). Finally, sub-clustering of the myeloid compartment revealed a mild shift in proportion within the barrier-associated macrophages and dendritic cell populations (Supplementary Figure 4E to G).

**Figure 4:**
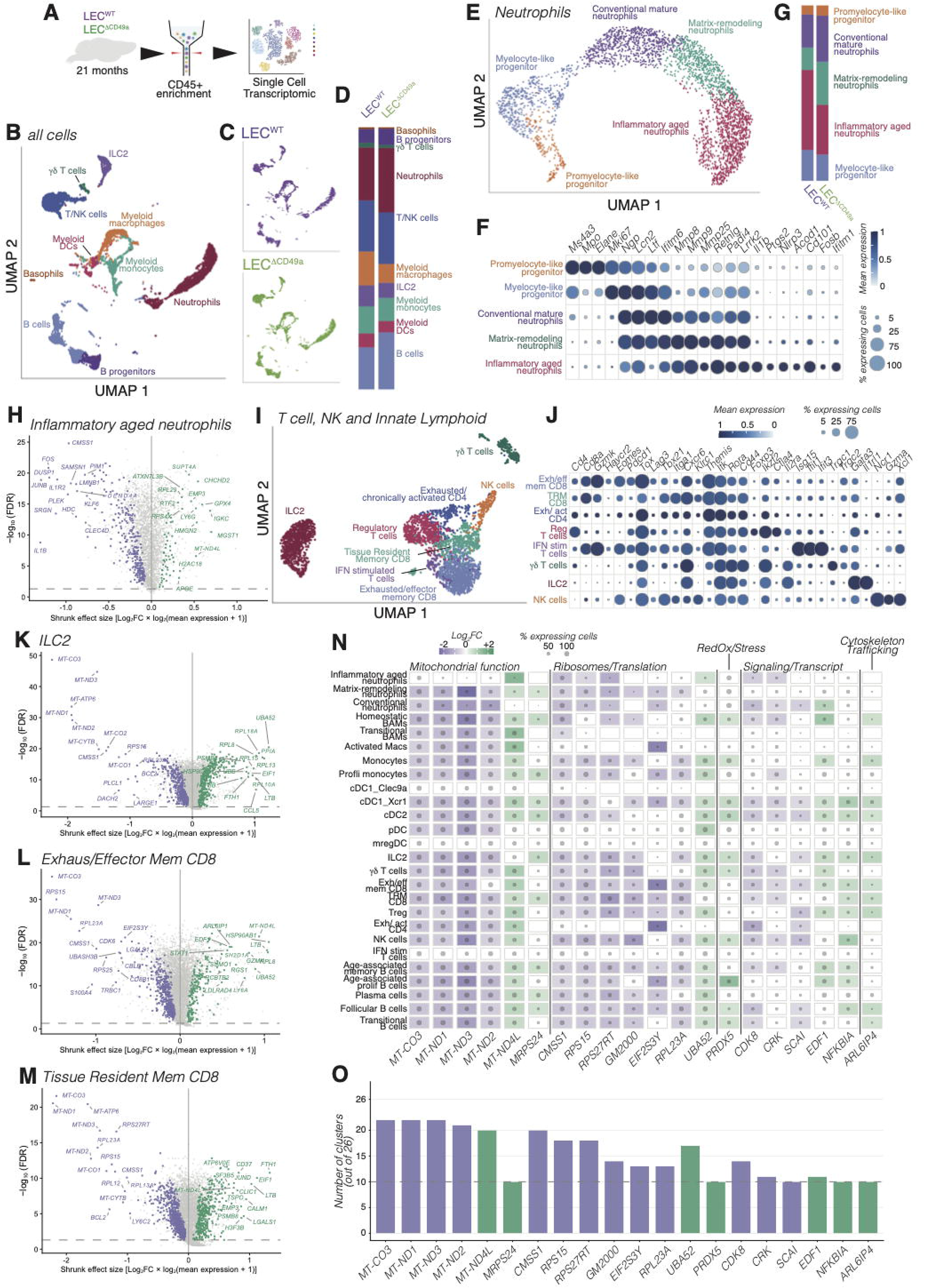
Loss of CD49a in LECs improves the aging phenotype of the dural immune compartment. A. Schematic of the experimental workflow. Cd45+ cells from aged LEC^WT^ and LEC^ΔCD49a^ dura were FACS sorted and sequenced using the 10x genomic platform. B. UMAP of all CD45+ cells colored by annotated cell type. C. UMAP split by genotype with LEC^WT^ (top, purple) and LEC^ΔCD49a^ (bottom, green) cells separately. D. Stacked bar plot of the relative proportion of each immune cell type in LEC^WT^ and LEC^ΔCD49a^ mice. E. UMAP of the neutrophil subset, colored by clusters. F. Dot plot of canonical marker genes defining each neutrophil subpopulation from (E). Dot color indicates mean expression; dot size indicated percentage of expressing cells. G. Stacked bar plot of the relative proportion of each neutrophil subpopulation in LEC^WT^ and LEC^ΔCD49a^ mice. H. Volcano plot of differentially expressed genes in the Inflammatory Aged Neutrophil cluster plotting shrunken effect size (Log2FC x log2(mean expression +1) against - log10(FDR). Genes upregulated LEC^ΔCD49a^ are shown in green, genes downregulated are shown in purple. I. UMAP of the T cells, NK and innate lymphoid cell subset, colored by annotated subtype. J. Dot plot of canonical marker genes defining each T/NK/ILC subpopulation from (I). Dot color indicates mean expression; dot size indicated percentage of expressing cells. K-M. Volcano plots of differentially expressed genes in ILC2 (K), exhausted/effector memory CD8 T cell (L) and tissue resident memory CD8 T cells (M), formatted as in (H) N. Heatmap of Log2FC for shared genes grouped by functional category across all annotated immune cell clusters. Color scale indicates Log2FC (purple, downregulated in LEC^ΔCD49a^, green upregulated in LEC^ΔCD49a^); dot size indicated percentage of expressing cells. O. Bar plot summarizing the number of immune cell clusters (out of 26 total) in which each gene from (N) is significantly differentially expressed, colored depending on sample upregulation (purple in LEC^WT^; green in LEC^ΔCD49a^).

To validate the immune composition predicted by the single-cell RNA sequencing, flow cytometry was performed on dural tissue of conditional CD49a KO mice and littermate controls (Supplementary Figure 5A). Specifically, CD11c expression on neutrophils was reduced in female, but not make, conditional KO mice (Supplementary Figure 5B and C). Interestingly, CD11c regulates ROS production by neutrophils (Koutsogiannaki et al., 2025; Hou et al., 2023), a process known to be increased in aging (Adrover et al., 2016). Moreover, we observed an increased in the percentage of CD62L+ CD44high CD4 T cells in the dura of conditional CD49a KO females, accompanied by a decreased proportion of CD62L+ CD4 T cells in the dCLNs (Supplementary Figure 5D to G). These changes were consistent with a shift away from a chronically activated, tissue-retained phenotype toward a more recirculation-competent memory-like phenotype. A previous study described an increase in regulatory T cells in the absence of the CCL21 receptor CCR7 (Da Mesquita et al., 2021a), however; we did not find the same change in our conditional KO (Supplementary Figure 5D). Lastly, we found an increase of the percentage of cDC1 dendritic cells (Cd11b-) with no change in the cDC2 (Cd11b+) population (Supplementary Figure 5H and I) in female conditional CD49a KO mice, suggesting genetic deletion of CD49a in LECs promotes remodeling of the DC compartment.

The recurrence of a mitochondrial/ribosomal signature across independent clusters prompted us to ask whether there was a compartment-wide, rather than cell-type-restricted response to the loss of CD49a in LECs. We therefore performed an unbiased cross-cluster analysis to identify genes that were significantly, differentially expressed in the same direction across all 26 independently defined clusters. This analysis revealed that core mitochondrial electron transport genes were significantly altered in a concordant manner across most clusters (20-22 out of 26). These changes were accompanied by a similarly broad changes in ribosomal and translation-associated genes, with more selective alterations in redox/stress response and cytoskeletal/trafficking programs (Figure 4N and O). Importantly, several observations supported that this shared transcriptional signature reflected a biological response to CD49a loss rather than a technical artifact. First, the signature was not universal, therefore inconsistent with a global technical confound. Secondly, genes of related function did not uniformly shift in the same direction; mitochondrial genes were consistently reduced in the conditional CD49a KO whereas ribosomal and translation-associated genes were increased in the same clusters, revealing a bidirectional, function-specific pattern inconsistent with technical artifact. These observations suggest that the loss of CD49a in LECs leads to a broadly shared metabolic shift in dural immune cells and improved cellular function. Overall, our data suggest that loss of CD49a in LECs promotes broad remodeling of the dural immune compartment in female mice, shifting immune populations away from an age-associated and chronically activated phenotype toward a more homeostatic state.

### Loss of lymphatic CD49a ameliorates cortical glial aging

Dural immune cells have been shown to influence the phenotype and function of surrounding brain parenchymal cells, including in the context of aging (Rustenhoven et al., 2023; Kaya et al., 2022; Filiano et al., 2016; Marin-Rodero et al., 2025). To determine if the changes in dural immunity may affect the brain of aged mice, we isolated the cortex and hippocampus of young and aged female conditional CD49a KO and WT littermates and performed targeted bulk RNA sequencing using the Nanostring platform (Figure 5A). We conducted gene set enrichment analysis (GSEA) using a combination of curated, literature-derived signatures of glial cell identity and reactivity (Supplementary Table 3), along with established Hallmark and ImmuneSigDB gene set collections. Analysis of the hippocampus did not show significant enrichment of any pathways. Similar to the single-cell RNA sequencing of the LECs, only one pathway was significantly enriched in the cortex of young conditional CD49a KO samples, suggesting a limited effect of the absence of CD49a in young mice (Supplementary Figure 6A and B). In the aged samples, however, GSEA analysis revealed significant negative enrichment for the aging microglia, disease-associated oligodendrocytes, and aging astrocytes signatures in the conditional CD49a KO mice, alongside reduced enrichment of IFNα and IFNγ response pathways (Figure 5B). These results suggest that LEC-specific loss of CD49a shifts cortical inflammatory and glial transcriptional programs away from an aging- and disease-associated states toward a more homeostatic profile. Examination of individual genes within these pathways demonstrated reduced expression of classical microglial aging and activation genes (*C1q*, *B2m*, *Lyz2*, *Ccl3*, *Cd68*), disease-associated oligodendrocytes markers (*Serpina3n*, *Tnfrsf12a* and *CD9*), and the aging-associated astrocytes markers (*Serpina3n* and *C3*) (Figure 5C).

**Figure 5:**
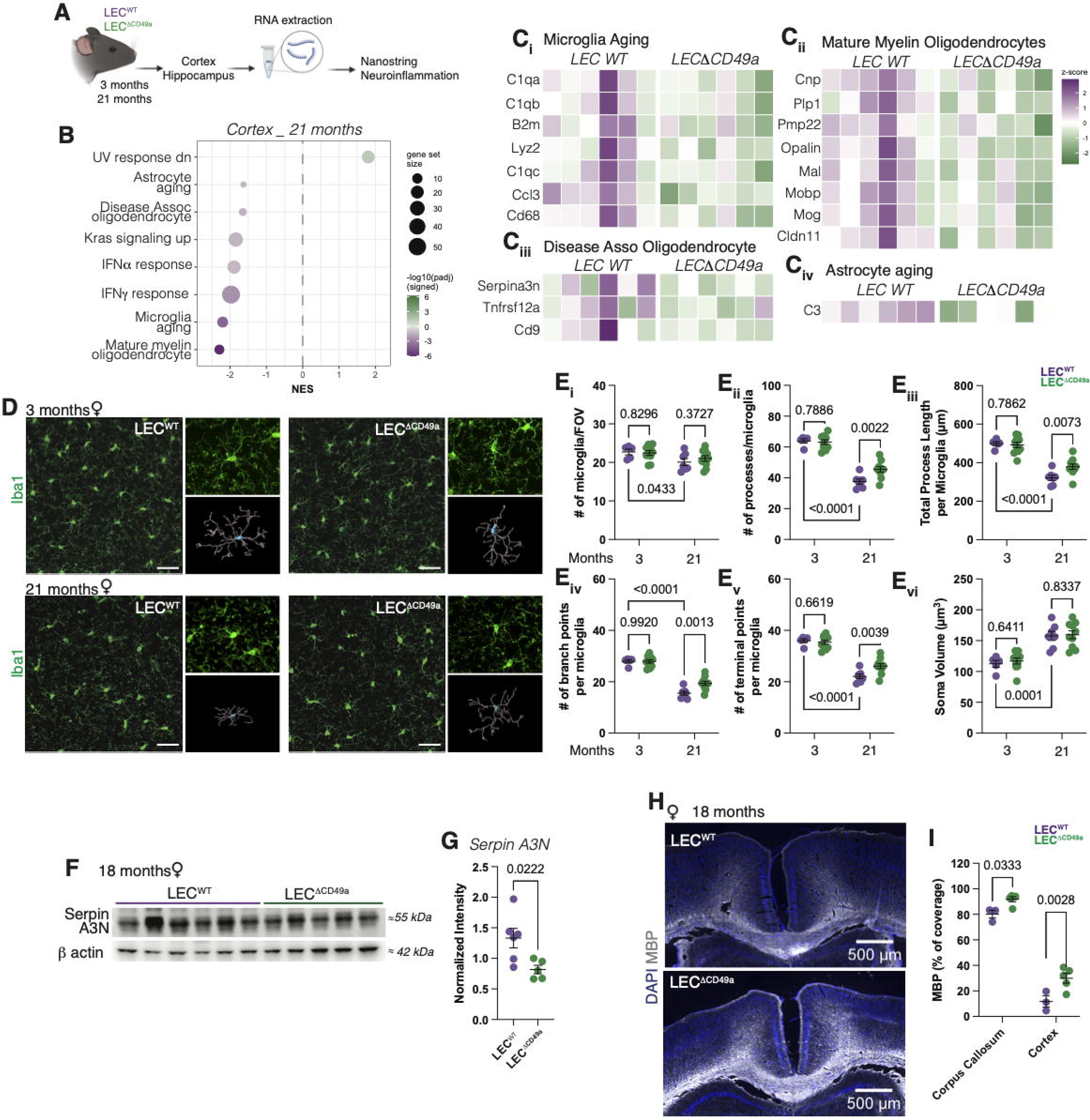
Improvement of aging phenotype across glial cells in absence of CD49a in LECs. A. Schematic of the experimental workflow. RNA was extracted from the cortex and hippocampus of young and aged LEC^WT^ and LEC^ΔCD49a^ mice. RNA abundances were measured using the Neuroinflammation panel of the Nanostring platform. B. Gene set enrichment analysis (GSEA) of sequenced genes comparing cortex from LEC^ΔCD49a^ to LEC^WT^ mice. Dot plot shows normalized enrichment score (NES) for each significantly enriched gene set. Dot size represented gene set size. C. Heatmaps of the z-scored expression for individual genes within gene sets identified in (B) comparing LEC^WT^ and LEC^ΔCD49a^ mice. Each column represents an individual biological replicate. D. Representative images of Iba1+ microglia in the cortex of LEC^WT^ and LEC^ΔCD49a^ mice at 3 months (top) and 21 months (bottom). For each genotype/age, the low-magnification field of view is shown on the left, with zoomed single microglia inset and its corresponding 3D-reconstructed skeletal trace used for morphometric analysis. E. Quantification of the number of microglia (i), number of processes per microglia (ii), total length of processes per microglial (iii), number of branch points per microglia (iv), number of terminal points per microglia (v) and microglial soma volume (vi) in young and aged LEC^WT^ and LEC^ΔCD49a^ mice. Mean ± s.e.m. Two-way ANOVA with Uncorrected Fisher’s LSD. F. Representative western blot panel of SerpinA3N and beta actin expression in the cortesx of LEC^WT^ and LEC^ΔCD49a^ at 18 months of age. G. Quantification of the normalized expression of SerpinA3N in the cortex of LEC^WT^ and LEC^ΔCD49a^ mice. Mean ± s.e.m. Welch’s t test. H. Representative images of MBP staining in the cortex and corpus callosum of aged LEC^WT^ and LEC^ΔCD49a^ mice. I. Quantification of the percentage of coverage of Mbp staining in the cortex and corpus callosum of LEC^WT^ and LEC^ΔCD49a^ mice. Mean ± s.e.m. Two-way ANOVA with Sidak’s multiple comparison test.

To determine whether these transcriptomic signatures were reflected in microglial morphology, we performed Iba1 immunostaining and 3D morphological reconstruction of cortical microglia in young and aged conditional CD49a KO mice and littermate controls. As expected (Hefendehl et al., 2014; Damani et al., 2011), aging in control mice was associated with a transition toward a less ramified, more amoeboid microglial morphology, characterized by reduced process number, process length, branch and terminal points, along with increased soma volume (Figure 5D and E). In aged conditional CD49a KO mice, this age-associated morphological remodeling was significantly attenuated compared to control mice, with increased process number, total process length, branch points, and terminal points (Figure 5D and E), while microglia density and soma volume remained unchanged. This effect was age- and sex-specific, as no morphological differences were observed in young mice of either genotype or in any microglia of male mice (Figure 5D and E, Supplementary Figure 6C). This microglial morphological improvement is consistent with the reduced aging-associated transcriptional signatures and suggest that CD49a loss preserves a more youthful microglial morphology in the aged cortex.

In addition to microglia, we also examined other glial cell population including oligodendrocytes. While CD49a loss did not alter the proportion of mature or immature oligodendrocyte populations in the cortex and corpus callosum (Supplementary Figure 6D and E), conditional CD49a KO mice female mice exhibited decreased cortical expression of SerpinA3N compared with controls (Figure 5F and G). SerpinA3N is a protein expressed by both disease-associated oligodendrocytes and astrocytes (Zhu et al., 2024), and its downregulation therefore illustrates a decrease in the pathological status of glial cells withing the brain with CD49a deletion in LECs. Furthermore, aging is associated with the loss of myelin in the cortex (Hill et al., 2018), prompting us to analyze myelination. We found a significant increase in MBP staining coverage in both the cortex and corpus callosum of female conditional CD49a KO mice compared to their WT littermates at 18 months of age (Figure 5H and I). Together, these findings demonstrate that LEC-specific CD49a loss attenuates aging-associated changes within the oligodendrocyte lineage and is associated with the preservation of myelin. Overall, our data suggest that loss of CD49a in LECs promotes a more homeostatic glial state in the aged cortex.

### Improvement of age-associated social and cognitive decline in absence of CD49a in LECs

Given the impact of CD49a loss in LECs on the dural immune compartment and the cortical glial cells, we next investigated whether these cellular changes translated into functional alterations in behavior. We first assessed open field and elevated plus maze performance in both male and female mice and saw no genotype difference in either young or aged mice (Supplementary Figure 7A to C). This indicates that any behavioral phenotype is not secondary to altered locomotion or anxiety-like behavior. In the 3-chambers social assay, at 3 months of age, both conditional KO and WT female littermates show a preference for the mouse over the object but by 18 months, this preference was lost in control mice while retained in conditional KO mice (Figure 6A). By 21 months, neither genotype exhibited a significant group level preference (Figure 6A). However, the sociability ratio showed a significant overall effect of genotype across ages and the proportion of mice classified as strongly sociable (discrimination index >1.5) was significantly higher in conditional CD49a KO mice compared to WT mice specifically at 21 months (Figure 6B and C). These effects were sex-specific, as male mice showed no genotype difference at either age (Supplementary Figure 7D and E).

**Figure 6:**
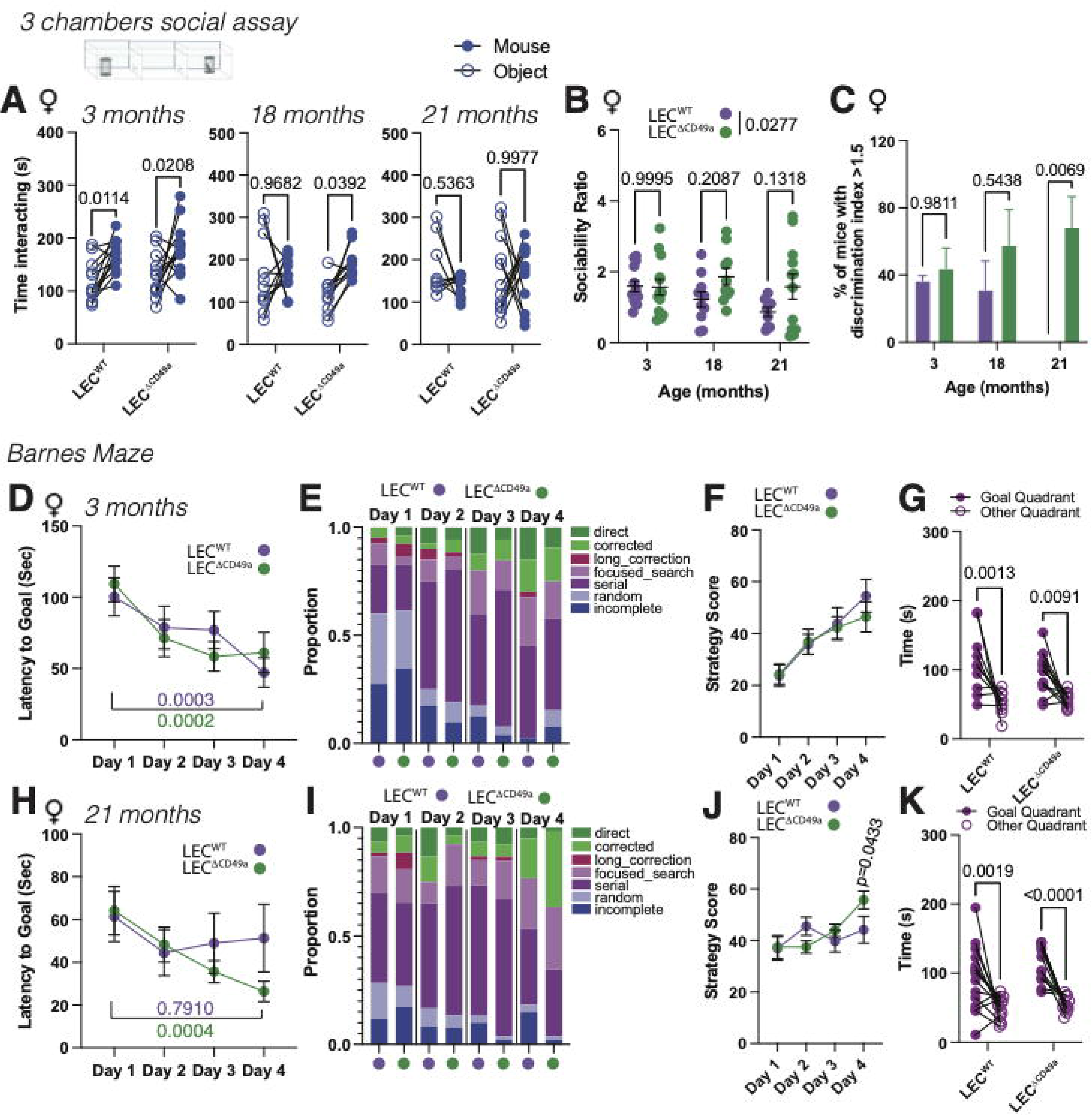
Age-associated behavioral improvement in mice lacking CD49a expression in LECs. A. Quantification of the time spent interacting with a mouse or inanimate object in the 3-chamber social assay at 3, 18 and 21 months of age comparing LEC^WT^ and LEC^ΔCD49a^ female mice. Mean ± s.e.m. Two-way ANOVA with Sidak’s multiple comparisons test. B. Quantification of the sociability ratio between LEC^WT^ and LEC^ΔCD49a^ female mice at 3, 18 and 21 months of age. Mean ± s.e.m. Two-way ANOVA with Sidak’s multiple comparisons test. C. Quantification of the percentage of mice with a discrimination index superior to 1.5 in LEC^WT^ and LEC^ΔCD49a^ female mice at 3, 18 and 21 months of age. Mean ± s.e.m. Two-way ANOVA with Sidak’s multiple comparisons test. D. Quantification of the latency to reach the goal hole during the training portion of the Barnes maze at 3 months of age. N=10/13 mice/group. Mean ± s.e.m. Two-way ANOVA with Sidak’s multiple comparisons test. E. Quantification of the proportion of strategy usage at 3 months of age. F. Quantification of the strategy score over the training days at 3 months of age. N=10/13 mice/group. Mean ± s.e.m. Two-way ANOVA with Sidak’s multiple comparisons test. G. Quantification of the time spent in the goal quadrant compared to the other quadrant during the test phase of the Barnes maze at 3 months of age. N=10/13 mice/group. Mean ± s.e.m. Two-way ANOVA with Sidak’s multiple comparisons test. H. Quantification of the latency to reach the goal hole during the training portion of the Barnes maze at 21 months of age. N=13/15 mice/group. Mean ± s.e.m. Two-way ANOVA with Sidak’s multiple comparisons test. I. Quantification of the proportion of strategy usage at 21 months of age. J. Quantification of the strategy score over the training days. At 21 months of age N=13/15 mice/group. Mean ± s.e.m. Two-way ANOVA with Sidak’s multiple comparisons test. K. Quantification of the time spent in the goal quadrant compared to the other quadrant during the test phase of the Barnes maze at 21 months of age. N=13/15 mice/group. Mean ± s.e.m. Two-way ANOVA with Sidak’s multiple comparisons test.

In the Barnes maze, 3 months-old mice of both genotypes demonstrated improved learning across training, with significantly reduced latency to reach the target hole from day 1 to day 4. This improvement was reflected by a comparable shift toward more efficient strategies, with increased use of goal-directed approaches and reduced random or incomplete searches across training days, with no differences in strategy scores between genotypes (Figure 6D to F, Supplementary Figure 7F). During testing, mice discriminated the goal quadrant compared to the other quadrants regardless of genotype (Figure 6G), suggesting intact memory retrieval. At 18 months, conditional CD49a KO and WT littermates showed comparable reductions in latency across training days (Supplementary Figure 7G). Despite similar learning curves, conditional CD49a KO mice demonstrated improved search strategies, with greater use of direct and corrected strategies and fewer incomplete or random strategies compared to WT (Supplementary Figure 7H), resulting in increased in strategy score in the conditional CD49a KO (Supplementary Figure 7I). Consistent with findings at 3 months of age, analysis of the test day showed no differences between genotype at 18 months (Supplementary Figure 7J). At 21 months of age, however, the behavioral phenotype was more pronounced, with conditional CD49a KO mice demonstrating a significant reduction in latency to reach the goal hole between day 1 and day 4 of training, whereas WT mice did not exhibit comparable improvement (Figure 6H). This enhanced learning was accompanied by increased use of efficient strategies and a significantly higher strategy score in conditional CD49a KO mice compared with control mice (Figure 6I and J). During the probe trial, both genotypes retained a significant preference for the goal quadrant, although this preference was modestly increased in the conditional KO mice (Figure 6K). As with the previous analysis, these observations were sex-specific, with male mice showing no statistical differences at any age (Supplementary Figure 7K to V). Together, these results demonstrate that CD49a loss in LECs preserves spatial learning capacity in aged female mice.

## Discussion

### Lymphatic aging is not defined by the loss of vessels alone

The concept of aging of the dural lymphatics has primarily revolved around the loss of lymphatic vessels and the associated decrease in bulk drainage of CSF and interstitial fluid (Da Mesquita et al., 2018b). Our data, along with others, indicates that changes in LEC functionality, independent of morphological alterations, also contribute to the aging of the dural lymphatics. Age-related impairment of CSF and macromolecule drainage has been recently shown to, at least partially, result from T-cell derived IFNγ acting directly on LECs to disrupt junctional integrity and increase paracellular permeability (Rustenhoven et al., 2023). Our data demonstrate that loss of immune cell drainage capacity similarly regulates aging of the dural lymphatics. Lymphatic-specific deletion of CD49a did not alter morphology or the bulk CSF drainage in aged mice, but selectively increased CCL21 expression by dural LECs and enhanced drainage of immune cell. Remarkably, this targeted improvement was sufficient to improve social and cognitive function. Together, these findings indicate that fluid drainage and immune drainage represent partially distinct cellular programs within the aging lymphatic endothelium. As a result, therapeutic strategies aimed solely at preventing lymphatic loss or promoting lymphatic growth in aged mice may only partially mitigate lymphatic-associated brain aging. Instead, targeting pathways that regulate specific lymphatic functions could provide a more efficient approach to improve age-associated behavioral decline.

Moreover, the functional role of CD49a in LECs appears to be confined to the aging, as deletion of CD49a produced no detectable phenotype in young mice. This age dependence may be reflective of changes in ECM-mediated signaling, as collagen IV composition, CD49a’s primary ligand (Kern et al., 1993), has been demonstrated to increase in the aged dura (Hitpass Romero et al., 2025). This observation also raises the question of what drives CD49a upregulation in aging. Notably, aging is accompanied by remodeling of the vascular and perivascular ECM (Ribeiro-Silva et al., 2021), characterized by collagen deposition, crosslinking, and basement membrane thickening (Santisteban and Iadecola, 2025; Hu et al., 2023). Since integrin expression is itself regulated by the composition and mechanical properties of the surrounding matrix (Yeh et al., 2017; Shih et al., 2011), the age-related induction of CD49a may reflect an adaptive response of LECs to a progressively collagen-rich dural microenvironment. Potential upstream drivers of CD49a induction include local pro-fibrotic mediators such as TGFβ, which has been shown to transcriptionally induce expression of other collagen-binding integrins including α11β1 (Lu et al., 2010; Talior-Volodarsky et al., 2015). TGFβ has also been recently demonstrated to impact dural fibrosis and lymphatic function (Hitpass Romero et al., 2025). Identifying the mechanisms that regulate CD49a upregulation in aging represents an important question to better understand the signals that drive lymphatic aging.

### Loss of CCL21 is a driver of dural aging

Our data support a model in which CD49a limits immune cell drainage by impairing the vesicular trafficking and secretory release of CCL21. However, our findings revealed a context-dependent effect of CD49a modulation on intracellular CCL21 levels with chronic CD49a activation increased CCL21 accumulation *in vitro*, whereas CD49a loss increased intracellular CCL21 in aged mice *in vivo*. We propose that this apparent discrepancy is linked to CCL21 demand. *In vitro*, in the absence of immune cells, CCL21 is released at a baseline level (Vaahtomeri et al., 2017; Johnson and Jackson, 2010). Therefore, the lack of vesicular transport and subsequent release of CCL21 leads to its accumulation within the cell. Contrastingly, *in vivo*, CCL21 release is regularly triggered by circulating immune cells (Russo et al., 2016; Weber et al., 2013) causing a constant recycling of the CCL21 pool. As aging progresses, increased CD49a expression begins to limit the rate of vesicle biogenesis and pool replenishment, ultimately decreasing the intracellular concentration of CCL21. This model is consistent with the coordinated downregulation of the SNARE-mediated exocytosis components - ORAI1, STX4, and VAMP5 - and the CCL21-vesicle markers RAB6 and RAB27b upon chronic CD49a activation. Future studies to directly measure the dynamics of CCL21 release downstream of CD49a stimulation are required to further refine the molecular link between CD49a and Ccl21 release.

Additionally, we identified a distinct mitochondrial phenotype associated with CD49a activation, characterized by altered representation of respiratory chain complexes and increased ROS production. Given that regulated vesicle trafficking and SNARE-mediated exocytosis are ATP-dependent processes (Jahn and Fasshauer, 2012; Cipriano et al., 2013), we hypothesize that the mitochondrial and trafficking phenotypes we describe are mechanistically linked with CD49a-driven mitochondrial reprogramming that limits the metabolic capacity required to sustain CCL21 release. Distinguishing whether mitochondrial dysfunction is upstream of, downstream of, or parallel to the trafficking defect would require metabolic perturbation in the context of CD49a gain- and loss-of-function.

Previous work has investigated meningeal immunity and CSF dynamics in CCR7-deficient mice (Da Mesquita et al., 2021a), the main receptor for CCL21. Global deletion of CCR7 leads to the accumulation of meningeal regulatory T cells, accompanied by impaired glymphatic influx and cognitive decline (Da Mesquita et al., 2021a). In contrast to these findings, we did not observe a change in regulatory T cell numbers with conditional deletion of CD49a in LECs, despite affecting CCL21 signaling. CCL19, the other chemokine that binds to CCR7, does not appear to depend on vesicular packaging for its release (Jørgensen et al., 2018; Bryce et al., 2016), suggesting it may remain available to sustain a degree of CCR7 signaling. Furthermore, a peripheral, non-LECs, source of CCL21 would similarly remain available (Manzo et al., 2007). This would predict a partial, rather than complete, phenocopy of CCR7 loss. As a result, our data establish that CD49a loss is sufficient to restore CCL21 levels and enhance immune drainage, but we have not directly established that these changes are responsible for the improved dural cell phenotype. Nevertheless, our data suggest that promoting the drainage of immune cells that became aberrantly retained in aged mice may improve the overall health of the dural compartment.

### Lymphatic endothelial cell reprogramming is an upstream driver of dural aging

A central question raised by our findings is how the loss of a single endothelial-intrinsic gene can drive broad remodeling of the dural immune compartment, affecting multiple cell types including neutrophils, B cells, CD4 T cells, and dendritic cells. In an egress model, restoring CCL21-dependent trafficking enhances chronically activated, retained, exhausted, or age-associated immune cells exit from the dura via the lymphatics and be replaced by recirculating or newly recruited cells, therefore promoting a more homeostatic dural immune environment. The loss of these cells may also simultaneously ameliorate the health of surrounding cells, similarly to how the clearance of senescent cells improves tissue function (Tizazu et al., 2025; Zhang et al., 2022), or how depletion of exhausted T cells improves the function of bystander cells (Minnie et al., 2022). Further, increased expression of immunoproteasome components, enhanced transcription and improvement in cellular transport may collectively result in secretion of extracellular vesicles (Wang et al., 2025; Zou et al., 2025; Ben-Nissan et al., 2022) by LECs that would benefit the local microenvironment (Trisko et al., 2022), including the different immune cell populations that reside primarily in proximity to the sinuses (Rustenhoven et al., 2021), where lymphatics are located (Louveau et al., 2015).

Moreover, the consequences of this immune remodeling appear to extend beyond the dura itself. Loss of CD49a in LECs is associated with attenuation of age-associated transcriptional profiles associated with cortical microglia, oligodendrocytes, and astrocytes, along with morphological changes in microglia. This is consistent with established literature demonstrating that aging CSF, dural, and infiltrating immune cells alter the status and function of parenchymal glial cells (Rustenhoven et al., 2023; Kaya et al., 2022). Notably, this rescue was selective for dampening reactive activity rather than altering cell number and was also regionally restricted, with no change observed in the hippocampus. This was paralleled at the behavioral level in which CD49a loss preferentially improved measures of learning acquisition and strategy efficiency, preserved social preference, and had modest effects on probe trial memory retention with aging. This pattern is consistent with a decrease in neuroinflammatory burden affecting circuits governing cortically mediated encoding behaviors. The regional pattern is also consistent with the influence of the close proximity between the dural/CSF compartment and the cortical regions (Iliff et al., 2012).

Further, the immune remodeling, glial rescue, and behavioral improvements associated with CD49a loss in LECs were consistently restricted to female mice, despite no sex preference in age-associated CD49a upregulation. The mechanistic basis for this sex specificity, whether hormonal, related to baseline sex differences in dural immune cells or demand for CCL21-dependent trafficking, or several additional factors, is not addressed by the present study and represents an important avenue for future investigation.

Finally, we note that CD49a was deleted from LECs throughout the entire adult lifespan of the animal, meaning our data demonstrates that CD49a loss prevents the emergence of age-associated lymphatic, immune, glial, and cognitive decline rather than preserves it once established. Whether inducing CD49a deletion in already-aged mice is sufficient to reverse the observed age-associated phenotypes is an important open question with direct bearing on the therapeutic potential of this pathway.

## Materials and Methods

### Mice

All mice used were on a C57Bl/6 background, bred in-house, and maintained under specific pathogen-free conditions on a 10/14hr light/dark cycle and ad libitum access to food and water. CD49a^flox/flox^ mice were generated by Cyagen from ES cells purchased from KOMP and crossed with Prox1^creERT2^ (gift from Dr Taija Makinen) to generate conditional CD49a lymphatic endothelial cells knockout mice, henceforth known as LEC^ΔCD49a^. All experiments were approved by the Institutional Animal Care and Use Committee of the Cleveland Clinic Research.

### Tamoxifen treatment

Tamoxifen (Sigma-Aldrich; T5648) was diluted into corn oil (Sigma-Aldrich; C8267) at a concentration of 20mg/ml. Tamoxifen was intraperitoneally injected into 6-8 weeks old mice and again at 17 months of age, including cre-negative littermates for 4 consecutive days at 80mg/kg body weight to induce conditional excision of CD49a from LECS.

### Behavioral testing and analysis

Behavioral tests were performed on mice at 3, 18 and 21 months of age for both male and female mice. Ethovision software (Noldus; version 13) was used to define the arenas, virtually generate the zones, track the mice during the behavioral tests, and for all subsequent analyses. All mice were allowed to habituate to the testing room for a minimum of 30 minutes prior to testing and returned to their home cage upon completion of the behavioral test.

#### Open field

Mice were placed in the center of a 45x45cm arena and allowed to freely explore for 10 minutes under white light at 100 lux. The thigmotaxis zone was designated to be the area 5cm from the perimeter of the arena and the center zone a 15x15cm area in the center of the arena. Each mouse underwent one trial.

#### Elevated Plus Maze

Mice were placed in the center of the maze that is composed of two enclosed 30cm long arms and two open 30cm long arm and allowed to feely explore the maze for 5 minutes under dim white light (25 lux). Each mouse underwent one trial.

#### 3 chamber social assay

Mice first underwent the habituation trial by being placed in the center chamber and allowed to freely explore all chambers for 10 minutes under white light at 100 lux. Immediately after the habituation trial, mice underwent the sociability trial where either a novel wildtype mouse (sex matched to the tested mouse) or inanimate object were placed in opposing chambers in an 8cm diameter wire cage. The test mouse was allowed to freely explore all the chambers for 10 minutes with the same lighting conditions as the previous trial.

#### Barnes Maze

The Barnes maze is a 100cm diameter disc with twenty 5cm holes around the periphery. For training, mice underwent four trials per day for four consecutive days with bright white light (>200lux). In the testing room, unique visual cues were placed on the walls (around 150cm from the edge of the maze). For each trial, mice were placed in the start box (at the center of the maze) for 15 seconds then allowed to freely explore the maze for up to 180 seconds. If the mouse did not find the goal box in the goal hole, the mouse was gently guided to the goal box where it remained for 60 seconds. Mice had at least 1h between trials. One the 5^th^ day, mice underwent the probe test, where the goal box was removed and the mouse was first placed in the start box for 15 seconds and then allowed to explore the Barnes maze for 5 minutes under bright light.

#### Tissue collection

Mice were euthanized by intraperitoneal injection of Beuthanasia (100mg/kg) and perfused with 0.1M of PBS (10ml/mouse). To isolate the brain and the meninges, skin was removed from the head, and the muscles were stripped from the skull. After section of the mandibles, and lateral cutting of the skull starting from the occipital hole, the brain was dislodged from the skull.

#### Immunohistochemistry and analysis

For staining of the dura, skull caps were harvested and post-fixed in 4% Normal Formalin Buffer (NFB; Thermo Fisher Scientific; SF100-4) for 24h. The dura was then stained in the skull cap and dissected afterwards and mounted onto slide. The lymph nodes were post-fixed in 4% NFB for 24h followed by 24h incubation in PBS with 30% sucrose. They were then frozen in OCT (Thermo Fisher Scientific; 23-730-571) and cryosectioned at 40µm sections. The brains were post-fixed in 4% NFB for 72h followed by 72h incubation in PBS with 30% sucrose then then frozen in OCT and cryosectioned at 15µm or 40µm sections. The tissues were washed 3x in 1X PBS and incubated for 1h in blocking solution (2% normal serum, 1% BSA (Equitech Bio; BAH70), 0.1% Triton-X-100 (Fisher Scientific; BP151), 0.05% Tween 20 (Fisher Scientific; BP337) and 0.1% Fc block – anti-CD16/CD32 (BioXCell; BE0307) in 1X PBS). After blocking, the tissues were incubated overnight in primary antibody solution (1% BSA, 0.5% Triton in 1% PBS + primary antibodies). The samples were then washed 3x in 1X PBS followed by 2h incubation in secondary antibody solution (1% BSA, 0.5% Triton in 1% PBS + secondary antibodies).

Following this step the sections were incubated in DAPI solution (1µg/ml in 1X PBS; Thermo Fisher; D1306) and washed 2x in 1X PBS. Whole mounts or sectioned tissues were mounted onto glass slides with Aqua-mount (Fisher Scientific; 14-39-5) and covered with glass cover slips. The images were acquired with Leica Stellaris confocal microscope at 10X of 40X magnification. The following primary antibodies were used: Rabbit polyclonal anti-Lyve-1 (1/800; ANGIOBIO Cat# 11-034), Rat monoclonal anti-Lyve-1 E660 (1/200; Thermo Scientific Cat# 50-0443-82), Rat monoclonal anti-CD3e biotin (1/500; Thermo Fisher Scientific Cat# 13-0032-82), Goat polyclonal anti-CCL21 (1/200; RCD Systems Cat# AF457), Goat polyclonal anti-Iba1 (1/500; Abcam Cat# ab5076), Rat monoclonal anti-MBP (1/200 Abcam Cat# ab7349), Mouse monoclonal anti-CC1 (1/200; Millipore Sigma; Cat# OP80), and Rabbit polyclonal anti-Olig2 (1/200; Millipore Sigma; AB9610). Fluorescent-labeled secondary antibodies or streptavidin targeting the primary antibody species or biotin were used for staining labeling except for already conjugated primary antibodies.

The morphology analysis of the meningeal lymphatics was performed with the Fiji ImageJ software (NIH). The whole length of the lymphatic vasculature along the transverse sinus was traced and measured with the freehand tool. Width was measured every 30-50µm of vessel segment along the transverse sinus with the measuring tool. OVA, Cd3e and Lyve-1 area coverage in the lymph nodes were measured on ImageJ by applying the same staining threshold across all images ang calculating the percentage of the LN area covered by the staining. CFSE-labeled T-cells were counted manually on ImageJ with the marking tool, and the density was calculated by dividing the number of cells / section area. Ccl21 in lymphatics was measured on ImageJ using the particles’ function. Microglia morphology was analyzed on IMARIS (OXFORD Instruments) using the filament tracing tool. MBP overage was assessed on ImageJ. An ROI was drawn to define the cortical and corpus callosum areas and the coverage was measured by applying the same staining threshold.

#### Flow cytometry

Meninges were digested for 15 min at 37°C with 1.4U/ml of Collagenase VIII (Sigma Aldrich; C2139) and 35U/ml of DNAse I (Sigma Aldrich; DN25) in complete media (DMEM (Gibco) with 2% FBS (Invitrogen), 1% L-glutamine (Gibco); 1% penicillin/streptomycin (Gibco); 1% sodium pyruvate (Gibco), 1% non-essential amino acid (Gibco), 1% essential amino acid (Gibco), and 1.5% HEPEC (Gibco)). The cell pellets were resuspended in ice-cold FACS Buffer (0.1M PBS; 1mM EDTA (Fisher Scientific) and 1% BSA (Equitech-Bio; BAH70); pH 7.4). Cells were stained for extracellular markers with antibodies at a 1:200 dilution. For intracellular staining, cells were fixed and permeabilized using the FoxP3 Buffer set (BD; 560409) following manufacturer’s instructions and cells were stained with antibodies at 1:200. The following antibodies were used : Rat anti-mouse CD45-Alexa 700 (BD; 560510); Syrian hamster anti-mouse podoplanin-PE (Thermo Fisher; 12-5381-82); Rat anti-mouse CD31-APC (Biolegend; 102509); Armenian hamster anti-mouse/rat CD49a-BV650 (BD; 740519); Rat anti-mouse CD11b-PE Cy7 (BD; 552850); Rat anti-mouse MHCII-PerCP Cy5.5 (Biolegend; 107626); Armenian hamster anti-mouse CD11c-APC (Biolegend; 117310); Rat anti-mouse F4/80-PE-CF596 (BD; 565613); Rat anti-mouse Ly6C-BV570 (Biolegend; 128030); Rat anti-mouse Ly6G (Biolegend; 127643); Rat anti-mouse Thy1.2-PerCP Cy5.5 (Biolegend; 140321); Mouse anti-mouse NK1.1-PE (Fisher Scientific; BDB561046); Armenian hamster anti-mouse TCRβ-BV711 (BD; 563135); Armenian hamster anti-mouse TCRγ/δ-PerCP Cy5.5 (Biolegend; 113117); Rat anti-mouse CD4-PE Cy7 (Fisher Scientific; BDB561099); Rat anti-mouse CD8a-V450 (BD; 560471); Mouse anti-mouse FoxP3-A647 (BD; 567462); Rat anti-mouse CD62L-A488 (Biolegend; 104419); Rat anti-mouse CD44-APC Cy7 (Biolegend; 103027). Cells were then fixed in 1% PFA in 0.1M pH7.4 PBS. Fluorescence data were collected with a Cytoflex (Beckman Coulter), then analyzed using FlowJo software (BD Life Sciences). Single cells were gated using the area and height of the forward scatter, then cells were selected for being live using the LIVE/DEAD Fixable Dead Cell Stain Kit per the manufacturer’s instruction (Invitrogen; L34957). An aliquot of unstained cells was counted using a Countess II cell counter (Thermo Fisher) using trypan blue to provide a cell counts from the samples.

#### LEC cell culture

Primary murine dermal lymphatic endothelial cells (LECs) were obtained commercially from Cell Biologics and grown according to provided protocols in Complete Mouse Endothelial Cell Medium (Cell Biologics, M1168). Briefly, LECs were isolated from the skin tissue of adult (Cell Biologics, C57-6064L) and aged (Cell Biologics, A57-6064L) (62-78 weeks) C57BL/6 pathogen-free laboratory mice and characterized by immunofluorescence for expression of VE-cadherin (AGED44, VE-cadherin Antibody, C-19, sc6458, Santa Cruz and AF1002, RCD Systems) or CD31/PECAM-1 (BD Pharmingen, 553370). Negative screening for bacteria, yeast, fungi, and mycoplasma was confirmed by Cell Biologics prior to cryopreservation. Following receipt, LECs were passaged, and cells up to P4 were used for all studies. qPCR analysis for LEC purity was conducted prior to initial use.

#### qPCR Validation of LECs

Total RNA was collected from untreated primary adult and aged LECs using an RNeasy Mini Kit (Qiagen, 74106) according to the manufacturer’s protocol and measured with a NanoDrop One spectrophotometer (ThermoFisher, ND-ONE-W). Total RNA was then normalized, treated with DNase I (ThermoFisher, 18068015), and reverse-transcribed to cDNA using a TaqMan Reverse Transcription kit (Applied Biosciences, N8080234). CDNA samples were then used for quantitative RT-PCR (qPCR) with SYBR Green reagents (Applied Biosystems, 4367659) and Taqman probes on a Quant Studio 6 Real-Time PCR system (ThermoFisher). Relative gene expression was quantified after normalizing to *Gapdh* expression. The following primer sequences were used: *Pecam* (F-CCAAAGCCAGTAGCATCATGGTC, R-GGATGGTGAAGTTGGCTACAGG), *Lyve1* (F-ACCAGGTAGAGTCAGCGCAGAA, R-CAGGACACCTTTGCCATTCTTCC), *Flt4* (Cat#4331182; Assay ID Mm01292604_m1), *Ccl21* (F-TCCCTACAGTATTGTCCGAGGC, R-ATCAGGTTCTGCACCCAGCCTT), *Pdpn* (F- ACAACCACAGGTGCTACTGGAG, R-GTTGCTGAGGTGGACAGTTCCT), *Ptprc* (F- CTTCAGTGGTCCCATTGTGGTG, R-TCAGACACCTCTGTCGCCTTAG), CCTCCCGTTATTGTGCAAGG). and *Pdgfra* (F- AGTGAGATCGAAGGCAGGCA, R-

#### CD49a activation time course

Adult and aged LECs were grown in gelatin-coated (Millipore Sigma, ES-006-B) tissue culture plates until reaching 90% confluency in Complete Mouse Endothelial Cell Medium. Cells were then synchronized in serum-free medium overnight. Following synchronization, cells were stimulated with 2 µg of an anti-CD49a antibody (BD Pharmingen, 555000) or a hamster IgG control (BD Pharmingen, 553961) for a period of 30 min, 1 h, or 4 h. Samples were then harvested using Accutase (MP Biomedicals, 091000449) and processed for Western blot.

#### Chronic CD49a activation

Adult and aged LECs were cultured in gelatin-coated tissue culture plates to 70% confluency in Complete Mouse Endothelial Cell Medium. After reaching the appropriate confluency, cells were stimulated with 2 µg of anti-CD49a antibody or a hamster IgG control. Cells were restimulated every 48 h with a 50% media change for a period of 7-8 days. Samples werehen harvested using Accutase and processed for Western blot.

#### Western Blot

Cultured LECs were harvested with Accutase and homogenized in RIPA lysis buffer (50 mM Tris-HCl pH = 7.4, 150 mM NaCl, 1 mM EDTA, 1% Triton X-100, and 1:100 proteinase/phosphatase inhibitor). Samples were briefly triturated, vortexed for 3 s, and incubated on ice for 30 min. For complete lysis, samples were frozen on dry ice for 3 min and thawed in wet ice for 15 min. A total of 3 freeze-thaw cycles were conducted. Samples were then centrifuged at 13,000 x *g* for 30 min, and supernatants were collected for immunoblotting. Protein levels were quantified using a Pierce BCA kit (ThermoFisher, 23227) and normalized. 4x Laemmli sample buffer (BioRad, 1610747) and b-mercaptoethanol (Sigma-Aldrich, M6250-100mL) were added to normalized samples. 10-20 μg of total protein lysate was resolved on Novex WedgeWell 4–20% Tris-Glycine gels (ThermoFisher, XP04202BOX) and transferred to PVDF membranes. The membranes were blocked with 3% bovine serum albumin (BSA) in Tris-buffered saline, 0.1% Tween 20 (TBST) for 1 h at room temperature and incubated overnight at 4°C with primary antibody. Membranes were blotted with primary antibodies for CCL21 (RCD Systems, MAB457), ORAI1 (Proteintech, 28411-1-AP), RAB6 (Cell Signaling, D37C7), Syntaxin 4 (Abcam, 184545), RAB27b (Proteintech, 13412-1-AP), VAMP5 (Proteintech, 11822-1-AP), TOM20 (Millipore Sigma, MABT166), VDAC1 (Abcam, N152B/23), b-actin (Invitrogen, MA5-15739), Total OXPHOS complexes (Abcam, STN-19467), phosho-FAK (Y397) (Cell Signaling, 3283S), total FAK (Cell Signaling, 3285S), phospho-SRC family (Y416) (Cell Signaling, 2101S), total SRC (Cell Signaling, 2108S), and CD49a (Cell Signaling, 71747S). The following day, membranes were washed with TBST and incubated with HRP-conjugated or fluorescently conjugated secondary antibodies for 1 h at room temperature. Membranes were then washed with TBST and imaged using the ChemiDoc MP imaging system (Bio-Rad) after activation with ECL substrate (Bio-Rad) or on an ODYSSEY CLx LI-COR system using Image Studio v5.2. Protein intensities were quantified using Image J software (v1.52a) and normalized to loading controls.

For tissue, Isolated cortices from whole brain flash frozen tissues. Once isolated, samples were homogenized in 10mL of chilled RIPA Lysis Buffer (Cell Signaling Technology, 1:100 protease/phosphatase inhibitor; 9806S) per gram of tissue. Samples were then centrifuged at 10,000g for 10 min and supernatants were collected for immunoblotting. Total protein concentration was determined using a Pierce BCA kit (ThermoFisher, 23225). Laemmli sample buffer (Bio-Rad; 1610747) and 2-mercaptoethanol (Sigma-Aldrich; M6250) were added to normalized samples. 15ug of total protein lysate was loaded on 4-20% Tris-Glycine Plus WedgeWell Gels (ThermoFisher, XP04205BOX). Following electrophoresis (117 V, 100 minutes), proteins were transferred (90V, 2 h) to nitrocellulose membranes (Bio-Rad, 1620115). The membranes were blocked with 3% bovine serum albumin in Tris-buffered saline and 0.1% Tween-20 (TBST) for 1 h at room temperature. Membranes were incubated overnight at 4° C with primary antibody. Membranes were blotted with primary antibodies for Serpin A3N (1:200; RCD Systems, AF4709). The following day, membranes were washed with TBST three times and incubated with horseradish peroxidase–conjugated antibodies (1:5000; Rat, Invitrogen, 31470) for 1 h at room temperature.

Membranes were then washed with TBST three times before activation with Clarity Max Western ECL Substrate (Bio-Rad; 1705062) and visualization on the ChemiDoc MP Imaging System (Bio-Rad). Membranes were stripped with Restore Western Blot Stripping Buffer (ThermoFisher Scientific; 2105) and then incubated for 1 h at room temperature with primary antibody the β-actin (1:1000; Cell Signaling Technology; 58169S). Following incubation, membranes were washed with TBST three times and incubated with horseradish peroxidase-conjugated antibodies (1:5000, Invitrogen, 31430) for 30 minutes at room temperature. Membranes were then washed with TBST three times before activation with Clarity Max Western ECL Substrate and visualization on the ChemiDoc MP Imaging System.

#### MitoSox staining

Adult and aged LECs were chronically stimulated with 2µg of anti-CD49a antibody or a hamster IgG control as previously described. Following treatment, cells were assessed for mitochondrial superoxide production using Mitosox. Media was aspirated and cells were incubated in 1mL of pre-warmed Hanks’ Balanced Salt Solution (HBSS) containing 500nm Mitosox Red (Invitrogen, M36008) for 30min at 37°C, 5% CO2, and protected from light. Following staining, cells were washed three times with 1mL of pre-warmed HBSS. Samples were then harvested using Accutase and processed for flow cytometry. Cell viability was assessed using a Near IR (876) fluorescent dye (Invitrogen, L34981A).

#### Cell isolation and single cell sequencing analysis

The cell preparation for the single cell RNA sequencing is the same as for regular flow cytometry with the addition of translation inhibitors (Actinomycin D (5µg/ml; Sigma Aldrich; A1410); Tripolide (10µM; Sigma Aldrich; T3652); Anisomycin (27.1µg/ml; Sigma Aldrich; A9789) to limit generation of digestion mediated artefact. Cells were stained with Rat anti-mouse CD45-A488 (Biolegend; 103121); Rat anti-mouse CD31-A647 (Biolegend; 102415) and Syrian hamster anti-mouse podoplanin-PE (Thermo Fisher; 12-5381-82). Prior to sorting, DAPI (5µg/ml in 1X PBS; Thermo Fisher; D1306) was added to the solution. Cells were sorted using FACS Aria III from the CCF flow cytometry core into BSA coated Eppendorf. CD31+ and CD45+ cells were then processed using the 10x Genomic Chromium platform (10x Genomics) following manufacturer’s instructions. Samples were sequenced using the Illumina NovaSeq using the S1 FlowCell.

Raw sequences reads were aligned to the mouse reference genome and gene expression matrices were generated using Cell Ranger (10x Genomics). Cell Ranger filtered genes were imported into BioTuring BBrowser for downstream analysis. Quality control filtering was applied to remove low-quality cells based on minimum gene count thresholds, maximum gene count thresholds (to exclude potential doublets), and percentage of reads mapping to mitochondrial genes. Genes detected in fewer than a minimum number of cells were also excluded from downstream analysis. Data from multiple samples were integrated to correct for batch effects using BioTuring’s built in integration. Normalized and scaled gene expression were used for principal component analysis (PCA), and the top principal components were used to construct a shared nearest neighbor (SNN) graph for unsupervised clustering. Cell clusters were visualized using Uniform Manifold Approximation and Projection (UMAP). Cell type identities were manually assigned to each cluster based on the expression of established marker genes for known dural cell population. Differentially expressed genes (DEGs) between male and female mice were identified within each cell type cluster using BioTuring BBrowwer.

For LEC analysis, DEGs were called using Venice statistical test and filtered based on log2 fold-change (above 0.25) and false discovery rate (inferior to 0.05). Cell type proportion were calculated with BioTuring BBrowser. DEGs were assigned to functional pathway categories based on annotated gene function (informed by g:Profiler pathway enrichment output together with manual curation). A custom R script (functional_network.R) rendered each gene as a node fill color mapped to Log2FC and node size mapped to -log10(FDR); a categorical overlay was used to indicated whether a gene reached significance in different comparison.

Age-associated DEGs were defined by comparing Young vs Old LEC^WT^. A gene was scored ‘rescued’ in the LEC^ΔCD49a^ condition if it was a significant age DEG whose direction of change was reversed (opposite sign, |Log2FC| > 0.5 in the reversing comparison) in Old LEC^ΔCD49a^ vs Old LEC^WT^. The reciprocal “reverse rescued” set was defined as genes significantly altered by CD49a deletion in aged LECs whose direction was likewise opposite to their behavior in the Young vs Old comparison. Set sizes and their intersection were computed in R and displayed as area-proportional Venn diagrams using the eulerr R package, with circle area scaled to gene-set size and the overlap representing genes concordant across both criteria.

For immune cells, DEGs were called using Venice statistical test and filtered based on log2 fold-change (above 0.5) and false discovery rate (inferior to 0.05). Marker and gene-expression dot plots were generated in R (ggplot2) following a shared template across cell-type/gene list panels: dot color encodes mean expression, and dot size encodes percent of expressing cells). Gene order was fixed manually per panel, and cluster/cell-type order followed the annotation scheme established for the corresponding UMAP.

Per cluster volcano plots were generated in R using a shrunk effect size on the x-axis, defined as shrunk effect size = Log2FC x log2(mean expression +1). This composite metric down-weights genes with low absolute expression so that highly significant but lowly expressed transcripts to not visually dominate the plot. Only well-expressed differentially expressed genes are prominently labeled. The y axis plots -log10(FDR). Genes were colored by direction of enrichment.

To identify transcriptional changes shared across meningeal immune populations, per-cluster DEGs tables were filtered at FDR < 0.05 and |Log2FC| > 0.05, and a gene was retained as part of the “shared signature” if it met this significance threshold, in the same direction, in at least 10 of the 26 clusters. Retained genes were manually grouped into functional categories for display. The resulting gene by cluster matrix was visualized as a heatmap generated in R and the number of clusters in which each shared gene reached significance was plotted as a bar chart.

#### Nanostring sequencing

RNA was extracted from cortex and hippocampus as previously described. 100ng of RNA per sample was used. The samples were analyzed with the nCounter Mouse Neuroinflammation (Cat# XT-CSO-MNROI1-12) using the nCounter analysis system according to manufacturer’s recommendation. The Bioturing SmartBulk platform was used to normalize the data using the nCounter pipeline, and for analysis.

Gene set enrichment analysis (GSEA) was used as the primary statistical framework. Genes were ranked for each comparison by a signed Wald statistic (log2 fold change divided by its standard error), reconstructed from the exported differential expression table. Preranked GSEA was performed using the fgsea R package (v1.39.4) with the fgseaMultilevel algorithm (minimum gene set size = 5 for MSigDB collections, 3 for the curated cell-type signatures; maximum size= 500, eps=0; seed= 42 for reproducibility).

Three gene set collections were tested: (1) the Hallmark collection (50 gene sets) and (2) the C7 ImmuneSigDB collection (4,872 gene sets), both obtained for Mus musculus via the msigdbr R package (MSigDD v7.5.1) and (3) a set of curated glial cell type/state gene signatures assembled from the literature (Supplementary Table 3). Gene sets were restricted to genes on the Nanostring neuroinflammation panel. Pathways with Benjamin-Hochberg-adjusted p<0.05 were considered statistically significant.

For each gene set, fgsea reports a normalized enrichment score (NES). NES sign indicates the direction of the enrichment relative to the ranking statistic. NES magnitude reflects the strength of enrichment and was used to rank and visualize pathways alongside adjusted p-value.

#### Statistics

Sample sizes were chosen based on standard power calculations using similar published experiments. Statistical methods were not used to recalculate or predetermine sample sizes. Statistical test usage is described in each figure. The numerical value of the statistical test is displayed on the graphs. Statistical outliers were removed based on the Grubbs test with a significance level of 0.05. Animals with biological (hydrocephaly, runt) or experimental (skull crack/CSF leak at injection, fail to perform behavior test, noticeable head/neck tumor in aged mice) abnormality were excluded from analysis. Choice of test were decided based on number of mice per group and distribution. Statistical analysis was performed using Prism 10 (GraphPad Software Inc).

#### Lead Contact

Request for further information and resources should be directed to and will be fulfilled by the lead contact Antoine Louveau

#### Resource Availability

This study did not generate new unique reagents.

#### Data and code availability

The scRNA-seq data will be deposited in the Gene Expression Omnibus (GEO) and made available on the date of publication.

## Acknowledgments

We thank all the members of the Louveau lab and the members of the Neuroimmunology group, particularly of the Williams Lab of the Neuroscience department of the Cleveland Clinic Research for their valuable inputs during discussions of this work. We also thank the Flow Cytometry Core and the Genomic Core of the Cleveland Clinic Research for the cell sorting and sequencing of our data. This work was supported by grant from the National Institutes of Health (R21AG077370) to AL. Graphical representations were created using BioRender (https://app.biorender.com/)

## Contributions

Concept: AL; Investigation: NMF, RT, GAT, CND, AL; Formal analysis: NMF, RT, GAT, CND, HB, NA, LC, ADB, AL; Writing: RT, GAT, CND, AL.

## References

Adrover, J.M., J.A. Nicolás-Ávila, and A. Hidalgo. 2016. Aging: A Temporal Dimension for Neutrophils. Trends Immunol. 37:334–345. doi:10.1016/j.it.2016.03.005.

Alves de Lima, K., J. Rustenhoven, and J. Kipnis. 2020. Meningeal Immunity and Its Function in Maintenance of the Central Nervous System in Health and Disease. Annu Rev Immunol. 38:597–620. doi:10.1146/annurev-immunol-102319-103410.

Aspelund, A., S. Antila, S.T. Proulx, T.V. Karlsen, S. Karaman, M. Detmar, H. Wiig, and K. Alitalo. 2015. A dural lymphatic vascular system that drains brain interstitial fluid and macromolecules. J Exp Med. 212:991–999. doi:10.1084/jem.20142290.

Barrientos, R.M., M.M. Kitt, L.R. Watkins, and S.F. Maier. 2015. Neuroinflammation in the normal aging hippocampus. Neuroscience. 309:84–99. doi:10.1016/j.neuroscience.2015.03.007.

Baruch, K., A. Deczkowska, E. David, J.M. Castellano, O. Miller, A. Kertser, T. Berkutzki, Z. Barnett-Itzhaki, D. Bezalel, T. Wyss-Coray, I. Amit, and M. Schwartz. 2014. Aging. Aging-induced type I interferon response at the choroid plexus negatively affects brain function. Science. 346:89–93. doi:10.1126/science.1252945.

Bazigou, E., S. Xie, C. Chen, A. Weston, N. Miura, L. Sorokin, R. Adams, A.F. Muro, D. Sheppard, and T. Makinen. 2009. Integrin-alpha9 is required for fibronectin matrix assembly during lymphatic valve morphogenesis. Dev Cell. 17:175–186. doi:10.1016/j.devcel.2009.06.017.

Ben-Nissan, G., N. Katzir, M.G. Füzesi-Levi, and M. Sharon. 2022. Biology of the Extracellular Proteasome. Biomolecules. 12:619. doi:10.3390/biom12050619.

Bottani, E., R. Cerutti, M.E. Harbour, S. Ravaglia, S.A. Dogan, C. Giordano, I.M. Fearnley, G. D’Amati, C. Viscomi, E. Fernandez-Vizarra, and M. Zeviani. 2017. TTC19 Plays a Husbandry Role on UQCRFS1 Turnover in the Biogenesis of Mitochondrial Respiratory Complex III. Mol Cell. 67:96–105.e4. doi:10.1016/j.molcel.2017.06.001.

Bromley, S.K., H. Akbaba, V. Mani, R. Mora-Buch, A.Y. Chasse, A. Sama, and A.D. Luster. 2020. CD49a Regulates Cutaneous Resident Memory CD8+ T Cell Persistence and Response. Cell Rep. 32:108085. doi:10.1016/j.celrep.2020.108085.

Bromley, S.K., S.Y. Thomas, and A.D. Luster. 2005. Chemokine receptor CCR7 guides T cell exit from peripheral tissues and entry into afferent lymphatics. Nat Immunol. 6:895–901. doi:10.1038/ni1240.

Bryce, S.A., R.A.M. Wilson, E.M. Tiplady, D.L. Asquith, S.K. Bromley, A.D. Luster, G.J. Graham, and R.J.B. Nibbs. 2016. ACKR4 on Stromal Cells Scavenges CCL19 To Enable CCR7-Dependent Trafficking of APCs from Inflamed Skin to Lymph Nodes. J Immunol. 196:3341–3353. doi:10.4049/jimmunol.1501542.

Chen, G., J.P. Schell, J.A. Benitez, S. Petropoulos, M. Yilmaz, B. Reinius, Z. Alekseenko, L. Shi, E. Hedlund, F. Lanner, R. Sandberg, and Q. Deng. 2016. Single-cell analyses of X Chromosome inactivation dynamics and pluripotency during differentiation. Genome Res. 26:1342–1354. doi:10.1101/gr.201954.115.

Chen, H., C. Griffin, L. Xia, and R.S. Srinivasan. 2014. Molecular and cellular mechanisms of lymphatic vascular maturation. Microvasc Res. 96:16–22. doi:10.1016/j.mvr.2014.06.002.

Cipriano, D.J., J. Jung, S. Vivona, T.D. Fenn, A.T. Brunger, and Z. Bryant. 2013. Processive ATP-driven substrate disassembly by the N-ethylmaleimide-sensitive factor (NSF) molecular machine. J Biol Chem. 288:23436–23445. doi:10.1074/jbc.M113.476705.

Clarke, L.E., S.A. Liddelow, C. Chakraborty, A.E. Münch, M. Heiman, and B.A. Barres. 2018. Normal aging induces A1-like astrocyte reactivity. Proc Natl Acad Sci U S A. 115:E1896–E1905. doi:10.1073/pnas.1800165115.

Cochemé, H.M., and M.P. Murphy. 2008. Complex I is the major site of mitochondrial superoxide production by paraquat. J Biol Chem. 283:1786–1798. doi:10.1074/jbc.M708597200.

Da Mesquita, S., Z. Fu, and J. Kipnis. 2018a. The Meningeal Lymphatic System: A New Player in Neurophysiology. Neuron. 100:375–388. doi:10.1016/j.neuron.2018.09.022.

Da Mesquita, S., J. Herz, M. Wall, T. Dykstra, K.A. de Lima, G.T. Norris, N. Dabhi, T. Kennedy, W. Baker, and J. Kipnis. 2021a. Aging-associated deficit in CCR7 is linked to worsened glymphatic function, cognition, neuroinflammation, and β-amyloid pathology. Science Advances. 7:eabe4601. doi:10.1126/sciadv.abe4601.

Da Mesquita, S., A. Louveau, A. Vaccari, I. Smirnov, R.C. Cornelison, K.M. Kingsmore, C. Contarino, S. Onengut-Gumuscu, E. Farber, D. Raper, K.E. Viar, R.D. Powell, W. Baker, N. Dabhi, R. Bai, R. Cao, S. Hu, S.S. Rich, J.M. Munson, M.B. Lopes, C.C. Overall, S.T. Acton, and J. Kipnis. 2018b. Functional aspects of meningeal lymphatics in ageing and Alzheimer’s disease. Nature. 560:185–191. doi:10.1038/s41586-018-0368-8.

Da Mesquita, S., Z. Papadopoulos, T. Dykstra, L. Brase, F.G. Farias, M. Wall, H. Jiang, C.D. Kodira, K.A. de Lima, J. Herz, A. Louveau, D.H. Goldman, A.F. Salvador, S. Onengut-Gumuscu, E. Farber, N. Dabhi, T. Kennedy, M.G. Milam, W. Baker, I. Smirnov, S.S. Rich, Dominantly Inherited Alzheimer Network, B.A. Benitez, C.M. Karch, R.J. Perrin, M. Farlow, J.P. Chhatwal, D.M. Holtzman, C. Cruchaga, O. Harari, and J. Kipnis. 2021b. Meningeal lymphatics affect microglia responses and anti-Aβ immunotherapy. Nature. 593:255–260. doi:10.1038/s41586-021-03489-0.

Dai, D., S. Gu, X. Han, H. Ding, Y. Jiang, X. Zhang, C. Yao, S. Hong, J. Zhang, Y. Shen, G. Hou, B. Qu, H. Zhou, Y. Qin, Y. He, J. Ma, Z. Yin, Z. Ye, J. Qian, Q. Jiang, L. Wu, Q. Guo, S. Chen, C. Huang, L.C. Kottyan, M.T. Weirauch, C.G. Vinuesa, and N. Shen. 2024. The transcription factor ZEB2 drives the formation of age-associated B cells. Science. 383:413–421. doi:10.1126/science.adf8531.

Damani, M.R., L. Zhao, A.M. Fontainhas, J. Amaral, R.N. Fariss, and W.T. Wong. 2011. Age-related alterations in the dynamic behavior of microglia. Aging Cell. 10:263–276. doi:10.1111/j.1474-9726.2010.00660.x.

D’Angelo, L., E. Astro, M. De Luise, I. Kurelac, N. Umesh-Ganesh, S. Ding, I.M. Fearnley, G. Gasparre, M. Zeviani, A.M. Porcelli, E. Fernandez-Vizarra, and L. Iommarini. 2021. NDUFS3 depletion permits complex I maturation and reveals TMEM126A/OPA7 as an assembly factor binding the ND4-module intermediate. Cell Rep. 35:109002. doi:10.1016/j.celrep.2021.109002.

Darshi, M., V.L. Mendiola, M.R. Mackey, A.N. Murphy, A. Koller, G.A. Perkins, M.H. Ellisman, and S.S. Taylor. 2011. ChChd3, an inner mitochondrial membrane protein, is essential for maintaining crista integrity and mitochondrial function. J Biol Chem. 286:2918–2932. doi:10.1074/jbc.M110.171975.

Davis, G.E. 1992. Affinity of integrins for damaged extracellular matrix: alpha v beta 3 binds to denatured collagen type I through RGD sites. Biochem Biophys Res Commun. 182:1025–1031. doi:10.1016/0006-291x(92)91834-d.

Deczkowska, A., H. Keren-Shaul, A. Weiner, M. Colonna, M. Schwartz, and I. Amit. 2018. Disease-Associated Microglia: A Universal Immune Sensor of Neurodegeneration. Cell. 173:1073–1081. doi:10.1016/j.cell.2018.05.003.

Dulken, B.W., M.T. Buckley, P. Navarro Negredo, N. Saligrama, R. Cayrol, D.S. Leeman, B.M. George, S.C. Boutet, K. Hebestreit, J.V. Pluvinage, T. Wyss-Coray, I.L. Weissman, H. Vogel, M.M. Davis, and A. Brunet. 2019. Single-cell analysis reveals T cell infiltration in old neurogenic niches. Nature. 571:205–210. doi:10.1038/s41586-019-1362-5.

Filiano, A.J., Y. Xu, N.J. Tustison, R.L. Marsh, W. Baker, I. Smirnov, C.C. Overall, S.P. Gadani, S.D. Turner, Z. Weng, S.N. Peerzade, H. Chen, K.S. Lee, M.M. Scott, M.P. Beenhakker, V. Litvak, and J. Kipnis. 2016. Unexpected role of interferon-γ in regulating neuronal connectivity and social behaviour. Nature. 535:425–429. doi:10.1038/nature18626.

Förster, R., A.C. Davalos-Misslitz, and A. Rot. 2008. CCR7 and its ligands: balancing immunity and tolerance. Nat Rev Immunol. 8:362–371. doi:10.1038/nri2297.

Franceschi, C., and J. Campisi. 2014. Chronic inflammation (inflammaging) and its potential contribution to age-associated diseases. J Gerontol A Biol Sci Med Sci. 69 Suppl 1:S4–9. doi:10.1093/gerona/glu057.

Galeeva, A., E. Treuter, S. Tomarev, and M. Pelto-Huikko. 2007. A prospero-related homeobox gene Prox-1 is expressed during postnatal brain development as well as in the adult rodent brain. Neuroscience. 146:604–616. doi:10.1016/j.neuroscience.2007.02.002.

Gartlan, K.H., J.L. Wee, M.C. Demaria, R. Nastovska, T.M. Chang, E.L. Jones, V. Apostolopoulos, G.A. Pietersz, M.J. Hickey, A.B. van Spriel, and M.D. Wright. 2013. Tetraspanin CD37 contributes to the initiation of cellular immunity by promoting dendritic cell migration. Eur J Immunol. 43:1208–1219. doi:10.1002/eji.201242730.

Ghezzi, D., P. Arzuffi, M. Zordan, C. Da Re, C. Lamperti, C. Benna, P. D’Adamo, D. Diodato, R. Costa, C. Mariotti, G. Uziel, C. Smiderle, and M. Zeviani. 2011. Mutations in TTC19 cause mitochondrial complex III deficiency and neurological impairment in humans and flies. Nat Genet. 43:259–263. doi:10.1038/ng.761.

Guan, S., L. Zhao, and R. Peng. 2022. Mitochondrial Respiratory Chain Supercomplexes: From Structure to Function. Int J Mol Sci. 23:13880. doi:10.3390/ijms232213880.

Hefendehl, J.K., J.J. Neher, R.B. Sühs, S. Kohsaka, A. Skodras, and M. Jucker. 2014. Homeostatic and injury-induced microglia behavior in the aging brain. Aging Cell. 13:60–69. doi:10.1111/acel.12149.

Hill, R.A., A.M. Li, and J. Grutzendler. 2018. Lifelong cortical myelin plasticity and age-related degeneration in the live mammalian brain. Nat Neurosci. 21:683–695. doi:10.1038/s41593-018-0120-6.

Hitpass Romero, K., T.J. Stevenson, L.C.D. Smyth, B. Watkin, S.J.C. McCullough, L. Vinnell, A.M. Smith, P. Schweder, J.A. Correia, J. Kipnis, M. Dragunow, and J. Rustenhoven. 2025a. Age-related meningeal extracellular matrix remodeling compromises CNS lymphatic function. J Neuroinflammation. 22:109. doi:10.1186/s12974-025-03436-0.

Hitpass Romero, K., T.J. Stevenson, L.C.D. Smyth, B. Watkin, S.J.C. McCullough, L. Vinnell, A.M. Smith, P. Schweder, J.A. Correia, J. Kipnis, M. Dragunow, and J. Rustenhoven. 2025b. Age-related meningeal extracellular matrix remodeling compromises CNS lymphatic function. J Neuroinflammation. 22:109. doi:10.1186/s12974-025-03436-0.

Hong, Y.-K., B. Lange-Asschenfeldt, P. Velasco, S. Hirakawa, R. Kunstfeld, L.F. Brown, P. Bohlen, D.R. Senger, and M. Detmar. 2004. VEGF-A promotes tissue repair-associated lymphatic vessel formation via VEGFR-2 and the alpha1beta1 and alpha2beta1 integrins. FASEB J. 18:1111–1113. doi:10.1096/fj.03-1179fje.

Hou, L., R.A. Voit, M. Shibamura-Fujiogi, S. Koutsogiannaki, Y. Li, Y. Chen, H. Luo, V.G. Sankaran, and K. Yuki. 2023. CD11c regulates neutrophil maturation. Blood Adv. 7:1312–1325. doi:10.1182/bloodadvances.2022007719.

Hu, Z., X. Deng, S. Zhou, C. Zhou, M. Shen, X. Gao, and Y. Huang. 2023. Pathogenic mechanisms and therapeutic implications of extracellular matrix remodelling in cerebral vasospasm. Fluids Barriers CNS. 20:81. doi:10.1186/s12987-023-00483-8.

Huang, G., H. Lu, A. Hao, D.C.H. Ng, S. Ponniah, K. Guo, C. Lufei, Q. Zeng, and X. Cao. 2004. GRIM-19, a cell death regulatory protein, is essential for assembly and function of mitochondrial complex I. Mol Cell Biol. 24:8447–8456. doi:10.1128/MCB.24.19.8447-8456.2004.

Iliff, J.J., M. Wang, Y. Liao, B.A. Plogg, W. Peng, G.A. Gundersen, H. Benveniste, G.E. Vates, R. Deane, S.A. Goldman, E.A. Nagelhus, and M. Nedergaard. 2012. A Paravascular Pathway Facilitates CSF Flow Through the Brain Parenchyma and the Clearance of Interstitial Solutes, Including Amyloid β. Science Translational Medicine. 4:147ra111–147ra111. doi:10.1126/scitranslmed.3003748.

Jahn, R., and D. Fasshauer. 2012. Molecular machines governing exocytosis of synaptic vesicles. Nature. 490:201–207. doi:10.1038/nature11320.

Johnson, L.A., and D.G. Jackson. 2010. Inflammation-induced secretion of CCL21 in lymphatic endothelium is a key regulator of integrin-mediated dendritic cell transmigration. Int Immunol. 22:839–849. doi:10.1093/intimm/dxq435.

Jørgensen, A.S., P.E. Adogamhe, J.M. Laufer, D.F. Legler, C.T. Veldkamp, M.M. Rosenkilde, and G.M. Hjortø. 2018. CCL19 with CCL21-tail displays enhanced glycosaminoglycan binding with retained chemotactic potency in dendritic cells. J Leukoc Biol. 104:401–411. doi:10.1002/JLB.2VMA0118-008R.

Jung, H.J., J.S. Shim, J. Lee, Y.M. Song, K.C. Park, S.H. Choi, N.D. Kim, J.H. Yoon, P.T. Mungai, P.T. Schumacker, and H.J. Kwon. 2010. Terpestacin inhibits tumor angiogenesis by targeting UQCRB of mitochondrial complex III and suppressing hypoxia-induced reactive oxygen species production and cellular oxygen sensing. J Biol Chem. 285:11584–11595. doi:10.1074/jbc.M109.087809.

Kataru, R.P., H.J. Park, J. Shin, J.E. Baik, A. Sarker, S. Brown, and B.J. Mehrara. 2022. Structural and Functional Changes in Aged Skin Lymphatic Vessels. Front Aging. 3:864860. doi:10.3389/fragi.2022.864860.

Kaya, T., N. Mattugini, L. Liu, H. Ji, L. Cantuti-Castelvetri, J. Wu, M. Schifferer, J. Groh, R. Martini, S. Besson-Girard, S. Kaji, A. Liesz, O. Gokce, and M. Simons. 2022. CD8+ T cells induce interferon-responsive oligodendrocytes and microglia in white matter aging. Nat Neurosci. 25:1446–1457. doi:10.1038/s41593-022-01183-6.

Kern, A., J. Eble, R. Golbik, and K. Kühn. 1993. Interaction of type IV collagen with the isolated integrins alpha 1 beta 1 and alpha 2 beta 1. Eur J Biochem. 215:151–159. doi:10.1111/j.1432-1033.1993.tb18017.x.

Knight, C.G., L.F. Morton, A.R. Peachey, D.S. Tuckwell, R.W. Farndale, and M.J. Barnes. 2000. The collagen-binding A-domains of integrins alpha(1)beta(1) and alpha(2)beta(1) recognize the same specific amino acid sequence, GFOGER, in native (triple-helical) collagens. J Biol Chem. 275:35–40. doi:10.1074/jbc.275.1.35.

Korin, B., T.L. Ben-Shaanan, M. Schiller, T. Dubovik, H. Azulay-Debby, N.T. Boshnak, T. Koren, and A. Rolls. 2017. High-dimensional, single-cell characterization of the brain’s immune compartment. Nat Neurosci. 20:1300–1309. doi:10.1038/nn.4610.

Koutsogiannaki, S., L. Hou, F. Alhamdan, M. Mastali, C. Murray, J. Van Eyk, K. Kunimura, and K. Yuki. 2025. DOCK2 as a novel CD11c ligand in neutrophils to regulate reactive oxygen species production. Front Immunol. 16:1692451. doi:10.3389/fimmu.2025.1692451.

Kriehuber, E., S. Breiteneder-Geleff, M. Groeger, A. Soleiman, S.F. Schoppmann, G. Stingl, D. Kerjaschki, and D. Maurer. 2001. Isolation and characterization of dermal lymphatic and blood endothelial cells reveal stable and functionally specialized cell lineages. J Exp Med. 194:797–808. doi:10.1084/jem.194.6.797.

Krishnarajah, S., F. Ingelfinger, E. Friebel, D. Cansever, A. Amorim, M. Andreadou, D. Bamert, G. Litscher, M. Lutz, M. Mayoux, S. Mundt, F. Ridder, C. Sparano, S.A. Stifter, C. Ulutekin, S. Unger, M. Vermeer, P. Zwicky, M. Greter, S. Tugues, D. De Feo, and B. Becher. 2022. Single-cell profiling of immune system alterations in lymphoid, barrier and solid tissues in aged mice. Nat Aging. 2:74–89. doi:10.1038/s43587-021-00148-x.

Kumar, J., R. Kumar, A. Kumar Singh, E.L. Tsakem, M. Kathania, M.J. Riese, A.L. Theiss, M.L. Davila, and K. Venuprasad. 2021. Deletion of Cbl-b inhibits CD8+ T-cell exhaustion and promotes CAR T-cell function. J Immunother Cancer. 9:e001688. doi:10.1136/jitc-2020-001688.

Labarta-Bajo, L., and N.J. Allen. 2025. Astrocytes in aging. Neuron. 113:109–126. doi:10.1016/j.neuron.2024.12.010.

Lavado, A., and G. Oliver. 2007. Prox1 expression patterns in the developing and adult murine brain. Dev Dyn. 236:518–524. doi:10.1002/dvdy.21024.

López-Otín, C., M.A. Blasco, L. Partridge, M. Serrano, and G. Kroemer. 2013. The hallmarks of aging. Cell. 153:1194–1217. doi:10.1016/j.cell.2013.05.039.

López-Otín, C., M.A. Blasco, L. Partridge, M. Serrano, and G. Kroemer. 2023. Hallmarks of aging: An expanding universe. Cell. 186:243–278. doi:10.1016/j.cell.2022.11.001.

Louveau, A., J. Herz, M.N. Alme, A.F. Salvador, M.Q. Dong, K.E. Viar, S.G. Herod, J. Knopp, J.C. Setliff, A.L. Lupi, S. Da Mesquita, E.L. Frost, A. Gaultier, T.H. Harris, R. Cao, S. Hu, J.R. Lukens, I. Smirnov, C.C. Overall, G. Oliver, and J. Kipnis. 2018. CNS lymphatic drainage and neuroinflammation are regulated by meningeal lymphatic vasculature. Nat Neurosci. 21:1380–1391. doi:10.1038/s41593-018-0227-9.

Louveau, A., I. Smirnov, T.J. Keyes, J.D. Eccles, S.J. Rouhani, J.D. Peske, N.C. Derecki, D. Castle, J.W. Mandell, K.S. Lee, T.H. Harris, and J. Kipnis. 2015. Structural and functional features of central nervous system lymphatic vessels. Nature. 523:337–341. doi:10.1038/nature14432.

Lu, N., S. Carracedo, J. Ranta, R. Heuchel, R. Soininen, and D. Gullberg. 2010. The human alpha11 integrin promoter drives fibroblast-restricted expression in vivo and is regulated by TGF-beta1 in a Smad- and Sp1-dependent manner. Matrix Biol. 29:166–176. doi:10.1016/j.matbio.2009.11.003.

Manzo, A., S. Bugatti, R. Caporali, R. Prevo, D.G. Jackson, M. Uguccioni, C.D. Buckley, C. Montecucco, and C. Pitzalis. 2007. CCL21 Expression Pattern of Human Secondary Lymphoid Organ Stroma Is Conserved in Inflammatory Lesions with Lymphoid Neogenesis. Am J Pathol. 171:1549–1562. doi:10.2353/ajpath.2007.061275.

Marin-Rodero, M., E. Cintado, A.J. Walker, T. Jayewickreme, F.A. Pinho-Ribeiro, Q. Richardson, R. Jackson, I.M. Chiu, C. Benoist, B. Stevens, J.L. Trejo, and D. Mathis. 2025. The meninges host a distinct compartment of regulatory T cells that preserves brain homeostasis. Sci Immunol. 10:eadu2910. doi:10.1126/sciimmunol.adu2910.

Masle-Farquhar, E., T.J. Peters, L.A. Miosge, I.A. Parish, C. Weigel, C.C. Oakes, J.H. Reed, and C.C. Goodnow. 2022. Uncontrolled CD21low age-associated and B1 B cell accumulation caused by failure of an EGR2/3 tolerance checkpoint. Cell Rep. 38:110259. doi:10.1016/j.celrep.2021.110259.

Mendrick, D.L., D.M. Kelly, S.S. duMont, and D.J. Sandstrom. 1995. Glomerular epithelial and mesangial cells differentially modulate the binding specificities of VLA-1 and VLA-2. Lab Invest. 72:367–375.

Mikhailik, A., B. Ford, J. Keller, Y. Chen, N. Nassar, and N. Carpino. 2007. A phosphatase activity of Sts-1 contributes to the suppression of TCR signaling. Mol Cell. 27:486–497. doi:10.1016/j.molcel.2007.06.015.

Minnie, S.A., O.G. Waltner, K.S. Ensbey, N.S. Nemychenkov, C.R. Schmidt, S.S. Bhise, S.RW. Legg, G. Campoy, L.D. Samson, R.D. Kuns, T. Zhou, J.D. Huck, S. Vuckovic, D. Zamora, A. Yeh, A. Spencer, M. Koyama, K.A. Markey, S.W. Lane, M. Boeckh, A.M. Ring, S.N. Furlan, and G.R. Hill. 2022. Depletion of exhausted alloreactive T cells enables targeting of stem-like memory T cells to generate tumor-specific immunity. Sci Immunol. 7:eabo3420. doi:10.1126/sciimmunol.abo3420.

Mrdjen, D., A. Pavlovic, F.J. Hartmann, B. Schreiner, S.G. Utz, B.P. Leung, I. Lelios, F.L. Heppner, J. Kipnis, D. Merkler, M. Greter, and B. Becher. 2018. High-Dimensional Single-Cell Mapping of Central Nervous System Immune Cells Reveals Distinct Myeloid Subsets in Health, Aging, and Disease. Immunity. 48:380–395.e6. doi:10.1016/j.immuni.2018.01.011.

Murphy, M.P. 2009. How mitochondria produce reactive oxygen species. Biochem J. 417:1–13. doi:10.1042/BJ20081386.

Nikolich-Žugich, J. 2018. The twilight of immunity: emerging concepts in aging of the immune system. Nat Immunol. 19:10–19. doi:10.1038/s41590-017-0006-x.

Norden, D.M., and J.P. Godbout. 2013. Review: microglia of the aged brain: primed to be activated and resistant to regulation. Neuropathol Appl Neurobiol. 39:19–34. doi:10.1111/j.1365-2990.2012.01306.x.

Novak, I., V. Kirkin, D.G. McEwan, J. Zhang, P. Wild, A. Rozenknop, V. Rogov, F. Löhr, D. Popovic, A. Occhipinti, A.S. Reichert, J. Terzic, V. Dötsch, P.A. Ney, and I. Dikic. 2010. Nix is a selective autophagy receptor for mitochondrial clearance. EMBO Rep. 11:45–51. doi:10.1038/embor.2009.256.

Olson, A.L., J.B. Knight, and J.E. Pessin. 1997. Syntaxin 4, VAMP2, and/or VAMP3/cellubrevin are functional target membrane and vesicle SNAP receptors for insulin-stimulated GLUT4 translocation in adipocytes. Mol Cell Biol. 17:2425–2435. doi:10.1128/MCB.17.5.2425.

Pang, X., X. He, Z. Qiu, H. Zhang, R. Xie, Z. Liu, Y. Gu, N. Zhao, Q. Xiang, and Y. Cui. 2023. Targeting integrin pathways: mechanisms and advances in therapy. Signal Transduct Target Ther. 8:1. doi:10.1038/s41392-022-01259-6.

Ramírez-Camacho, I., W.R. García-Niño, M. Flores-García, J. Pedraza-Chaverri, and C. Zazueta. 2020. Alteration of mitochondrial supercomplexes assembly in metabolic diseases. Biochim Biophys Acta Mol Basis Dis. 1866:165935. doi:10.1016/j.bbadis.2020.165935.

Randolph, G.J., S. Ivanov, B.H. Zinselmeyer, and J.P. Scallan. 2017. The Lymphatic System: Integral Roles in Immunity. Annu Rev Immunol. 35:31–52. doi:10.1146/annurev-immunol-041015-055354.

Ray, S.J., S.N. Franki, R.H. Pierce, S. Dimitrova, V. Koteliansky, A.G. Sprague, P.C. Doherty, A.R. de Fougerolles, and D.J. Topham. 2004. The collagen binding alpha1beta1 integrin VLA-1 regulates CD8 T cell-mediated immune protection against heterologous influenza infection. Immunity. 20:167–179. doi:10.1016/s1074-7613(04)00021-4.

Ribeiro-Silva, J.C., P. Nolasco, J.E. Krieger, and A.A. Miyakawa. 2021. Dynamic Crosstalk between Vascular Smooth Muscle Cells and the Aged Extracellular Matrix. Int J Mol Sci. 22:10175. doi:10.3390/ijms221810175.

Russo, E., A. Teijeira, K. Vaahtomeri, A.-H. Willrodt, J.S. Bloch, M. Nitschké, L. Santambrogio, D. Kerjaschki, M. Sixt, and C. Halin. 2016. Intralymphatic CCL21 Promotes Tissue Egress of Dendritic Cells through Afferent Lymphatic Vessels. Cell Rep. 14:1723–1734. doi:10.1016/j.celrep.2016.01.048.

Rustenhoven, J., A. Drieu, T. Mamuladze, K.A. de Lima, T. Dykstra, M. Wall, Z. Papadopoulos, M. Kanamori, A.F. Salvador, W. Baker, M. Lemieux, S. Da Mesquita, A. Cugurra, J. Fitzpatrick, S. Sviben, R. Kossina, P. Bayguinov, R.R. Townsend, Q. Zhang, P. Erdmann-Gilmore, I. Smirnov, M.-B. Lopes, J. Herz, and J. Kipnis. 2021. Functional characterization of the dural sinuses as a neuroimmune interface. Cell. 184:1000–1016.e27. doi:10.1016/j.cell.2020.12.040.

Rustenhoven, J., and J. Kipnis. 2022. Brain borders at the central stage of neuroimmunology. Nature. 612:417–429. doi:10.1038/s41586-022-05474-7.

Rustenhoven, J., G. Pavlou, S.E. Storck, T. Dykstra, S. Du, Z. Wan, D. Quintero, J.P. Scallan, I. Smirnov, R.D. Kamm, and J. Kipnis. 2023. Age-related alterations in meningeal immunity drive impaired CNS lymphatic drainage. J Exp Med. 220:e20221929. doi:10.1084/jem.20221929.

Santisteban, M.M., and C. Iadecola. 2025. The pathobiology of neurovascular aging. Neuron. 113:49–70. doi:10.1016/j.neuron.2024.12.014.

Sarris, M., J.-B. Masson, D. Maurin, L.M. Van der Aa, P. Boudinot, H. Lortat-Jacob, and P. Herbomel. 2012. Inflammatory chemokines direct and restrict leukocyte migration within live tissues as glycan-bound gradients. Curr Biol. 22:2375–2382. doi:10.1016/j.cub.2012.11.018.

Schweers, R.L., J. Zhang, M.S. Randall, M.R. Loyd, W. Li, F.C. Dorsey, M. Kundu, J.T. Opferman, J.L. Cleveland, J.L. Miller, and P.A. Ney. 2007. NIX is required for programmed mitochondrial clearance during reticulocyte maturation. Proc Natl Acad Sci U S A. 104:19500–19505. doi:10.1073/pnas.0708818104.

Segel, M., B. Neumann, M.F.E. Hill, I.P. Weber, C. Viscomi, C. Zhao, A. Young, C.C. Agley, A.J. Thompson, G.A. Gonzalez, A. Sharma, S. Holmqvist, D.H. Rowitch, K. Franze, R.J.M. Franklin, and K.J. Chalut. 2019. Niche stiffness underlies the ageing of central nervous system progenitor cells. Nature. 573:130–134. doi:10.1038/s41586-019-1484-9.

Seifert, U., L.P. Bialy, F. Ebstein, D. Bech-Otschir, A. Voigt, F. Schröter, T. Prozorovski, N. Lange, J. Steffen, M. Rieger, U. Kuckelkorn, O. Aktas, P.-M. Kloetzel, and E. Krüger. 2010a. Immunoproteasomes preserve protein homeostasis upon interferon-induced oxidative stress. Cell. 142:613–624. doi:10.1016/j.cell.2010.07.036.

Seifert, U., L.P. Bialy, F. Ebstein, D. Bech-Otschir, A. Voigt, F. Schröter, T. Prozorovski, N. Lange, J. Steffen, M. Rieger, U. Kuckelkorn, O. Aktas, P.-M. Kloetzel, and E. Krüger. 2010b. Immunoproteasomes preserve protein homeostasis upon interferon-induced oxidative stress. Cell. 142:613–624. doi:10.1016/j.cell.2010.07.036.

Sharifi, R., J.C. Sinclair, K.C. Gilmour, P.D. Arkwright, C. Kinnon, A.J. Thrasher, and H.B. Gaspar. 2004. SAP mediates specific cytotoxic T-cell functions in X-linked lymphoproliferative disease. Blood. 103:3821–3827. doi:10.1182/blood-2003-09-3359.

Shih, Y.-R.V., K.-F. Tseng, H.-Y. Lai, C.-H. Lin, and O.K. Lee. 2011. Matrix stiffness regulation of integrin-mediated mechanotransduction during osteogenic differentiation of human mesenchymal stem cells. J Bone Miner Res. 26:730–738. doi:10.1002/jbmr.278.

Sreejit, G., S.K. Nooti, R.M. Jaggers, B. Athmanathan, K. Ho Park, A. Al-Sharea, J. Johnson, A. Dahdah, M.K.S. Lee, J. Ma, A.J. Murphy, and P.R. Nagareddy. 2022. Retention of the NLRP3 Inflammasome-Primed Neutrophils in the Bone Marrow Is Essential for Myocardial Infarction-Induced Granulopoiesis. Circulation. 145:31–44. doi:10.1161/CIRCULATIONAHA.121.056019.

Summers deLuca, L., D. Ng, Y. Gao, M.E. Wortzman, T.H. Watts, and J.L. Gommerman. 2011. LTβR signaling in dendritic cells induces a type I IFN response that is required for optimal clonal expansion of CD8+ T cells. Proc Natl Acad Sci U S A. 108:2046–2051. doi:10.1073/pnas.1014188108.

Talior-Volodarsky, I., P.D. Arora, Y. Wang, C. Zeltz, K.A. Connelly, D. Gullberg, and C.A. McCulloch. 2015. Glycated Collagen Induces α11 Integrin Expression Through TGF-β2 and Smad3. J Cell Physiol. 230:327–336. doi:10.1002/jcp.24708.

Tizazu, A.M., E.A. Tessema, and O.NF. Cexus. 2025. Targeting immunosenesence promotes clearance of senescent cells. Immun Ageing. 22:35. doi:10.1186/s12979-025-00518-8.

Trisko, J., J. Fleck, S. Kau, J. Oesterreicher, and W. Holnthoner. 2022. Lymphatic and Blood Endothelial Extracellular Vesicles: A Story Yet to Be Written. Life (Basel). 12:654. doi:10.3390/life12050654.

Ushijima, M., T. Uruno, A. Nishikimi, F. Sanematsu, Y. Kamikaseda, K. Kunimura, D. Sakata, T. Okada, and Y. Fukui. 2018. The Rac Activator DOCK2 Mediates Plasma Cell Differentiation and IgG Antibody Production. Front Immunol. 9:243. doi:10.3389/fimmu.2018.00243.

Vaahtomeri, K., M. Brown, R. Hauschild, I. De Vries, A.F. Leithner, M. Mehling, W.A. Kaufmann, and M. Sixt. 2017. Locally Triggered Release of the Chemokine CCL21 Promotes Dendritic Cell Transmigration across Lymphatic Endothelia. Cell Rep. 19:902–909. doi:10.1016/j.celrep.2017.04.027.

Van Avondt, K., J.-K. Strecker, C. Tulotta, J. Minnerup, C. Schulz, and O. Soehnlein. 2023. Neutrophils in aging and aging-related pathologies. Immunol Rev. 314:357–375. doi:10.1111/imr.13153.

Van Bruggen, S., P.-A. Jarrot, E. Thomas, C.E. Sheehy, C.M.S. Silva, A.Y. Hsu, P. Cunin, P.A. Nigrovic, E.R. Gomes, H.R. Luo, C.M. Waterman, and D.D. Wagner. 2023. NLRP3 is essential for neutrophil polarization and chemotaxis in response to leukotriene B4 gradient. Proc Natl Acad Sci U S A. 120:e2303814120. doi:10.1073/pnas.2303814120.

Van Hove, H., L. Martens, I. Scheyltjens, K. De Vlaminck, A.R. Pombo Antunes, S. De Prijck, N. Vandamme, S. De Schepper, G. Van Isterdael, C.L. Scott, J. Aerts, G. Berx, G.E. Boeckxstaens, R.E. Vandenbroucke, L. Vereecke, D. Moechars, M. Guilliams, J.A. Van Ginderachter, Y. Saeys, and K. Movahedi. 2019. A single-cell atlas of mouse brain macrophages reveals unique transcriptional identities shaped by ontogeny and tissue environment. Nat Neurosci. 22:1021–1035. doi:10.1038/s41593-019-0393-4.

Vig, M., W.I. DeHaven, G.S. Bird, J.M. Billingsley, H. Wang, P.E. Rao, A.B. Hutchings, M.-H. Jouvin, J.W. Putney, and J.-P. Kinet. 2008. Defective mast cell effector functions in mice lacking the CRACM1 pore subunit of store-operated calcium release-activated calcium channels. Nat Immunol. 9:89–96. doi:10.1038/ni1550.

Vilchez, D., I. Morantte, Z. Liu, P.M. Douglas, C. Merkwirth, A.P.C. Rodrigues, G. Manning, and A. Dillin. 2012. RPN-6 determines C. elegans longevity under proteotoxic stress conditions. Nature. 489:263–268. doi:10.1038/nature11315.

Wang, W., S. Qiao, X. Kong, G. Zhang, and Z. Cai. 2025. The role of exosomes in immunopathology and potential therapeutic implications. Cell Mol Immunol. 22:975–995. doi:10.1038/s41423-025-01323-5.

Weber, M., R. Hauschild, J. Schwarz, C. Moussion, I. de Vries, D.F. Legler, S.A. Luther, T. Bollenbach, and M. Sixt. 2013. Interstitial dendritic cell guidance by haptotactic chemokine gradients. Science. 339:328–332. doi:10.1126/science.1228456.

Yeh, Y.-C., J.-Y. Ling, W.-C. Chen, H.-H. Lin, and M.-J. Tang. 2017. Mechanotransduction of matrix stiffness in regulation of focal adhesion size and number: reciprocal regulation of caveolin-1 and β1 integrin. Sci Rep. 7:15008. doi:10.1038/s41598-017-14932-6.

Zeng, Q., V.N. Subramaniam, S.H. Wong, B.L. Tang, R.G. Parton, S. Rea, D.E. James, and W. Hong. 1998. A Novel Synaptobrevin/VAMP Homologous Protein (VAMP5) Is Increased during In Vitro Myogenesis and Present in the Plasma Membrane. Mol Biol Cell. 9:2423–2437. doi:10.1091/mbc.9.9.2423.

Zhang, X., V.M. Pearsall, C.M. Carver, E.J. Atkinson, B.D.S. Clarkson, E.M. Grund, M. Baez-Faria, K.D. Pavelko, J.M. Kachergus, T.A. White, R.K. Johnson, C.S. Malo, A.M. Gonzalez-Suarez, K. Ayasoufi, K.O. Johnson, Z.P. Tritz, C.E. Fain, R.H. Khadka, M. Ogrodnik, D. Jurk, Y. Zhu, T. Tchkonia, A. Revzin, J.L. Kirkland, A.J. Johnson, C.L. Howe, E.A. Thompson, N.K. LeBrasseur, and M.J. Schafer. 2022. Rejuvenation of the aged brain immune cell landscape in mice through p16-positive senescent cell clearance. Nat Commun. 13:5671. doi:10.1038/s41467-022-33226-8.

Zhao, M.-Y., C.-Y. Ye, Y.-C. Liu, X.-M. Wang, J.-C. Fu, X.-Y. Liu, R. Zhu, Y.-Z. Li, and Q. Tian. 2025. Role of meningeal lymphatic vessels in brain homeostasis. Front Immunol. 16:1593630. doi:10.3389/fimmu.2025.1593630.

Zhou, Y., J. Cai, W. Zhang, X. Gong, S. Yan, K. Zhang, Z. Luo, J. Sun, Q. Jiang, and M. Lou. 2020. Impairment of the Glymphatic Pathway and Putative Meningeal Lymphatic Vessels in the Aging Human. Ann Neurol. 87:357–369. doi:10.1002/ana.25670.

Zhu, M., Z. Lan, J. Park, S. Gong, Y. Wang, and F. Guo. 2024. Regulation of CNS pathology by Serpina3n/SERPINA3: The knowns and the puzzles. Neuropathol Appl Neurobiol. 50:e12980. doi:10.1111/nan.12980.

Zolla, V., I.T. Nizamutdinova, B. Scharf, C.C. Clement, D. Maejima, T. Akl, T. Nagai, P. Luciani, J.-C. Leroux, C. Halin, S. Stukes, S. Tiwari, A. Casadevall, W.R. Jacobs, D. Entenberg, D.C. Zawieja, J. Condeelis, D.R. Fooksman, A.A. Gashev, and L. Santambrogio. 2015. Aging-related anatomical and biochemical changes in lymphatic collectors impair lymph transport, fluid homeostasis, and pathogen clearance. Aging Cell. 14:582–594. doi:10.1111/acel.12330.

Zou, Z., Y. Hao, Z. Tao, W. Ye, Z. Luo, X. Li, R. Li, K. Zheng, J. Xia, C. Guo, X. Zhang, and J. Wu. 2025. Current landscape of the immunoproteasome: implications for disease and therapy. Cell Death Discov. 11:406. doi:10.1038/s41420-025-02698-0.

