## Supplementary Material for "Lymphatic CD49a is a driver of meningeal immune aging and cognitive decline"

#### Supplementary Figures and Legends:

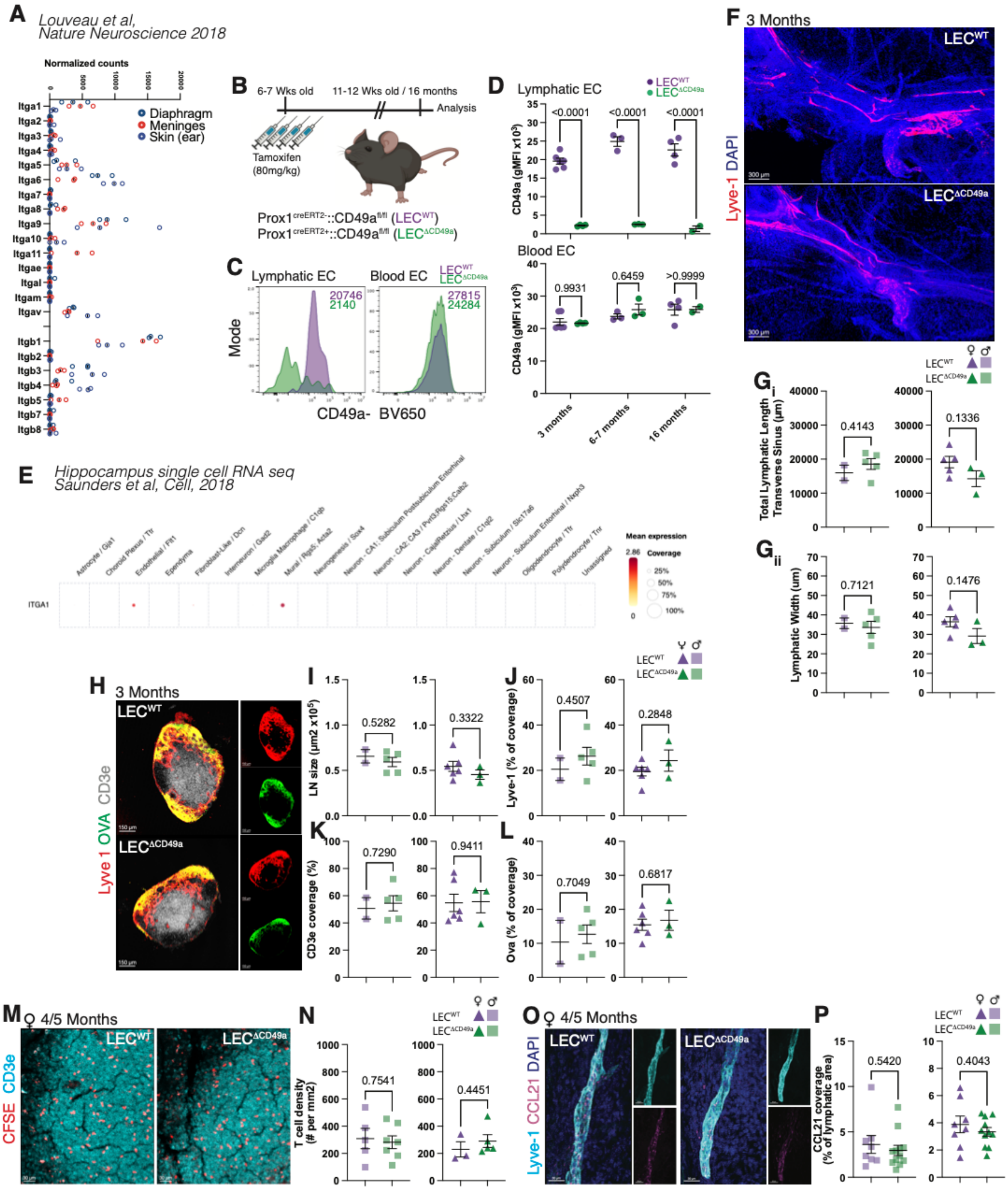

**Supplementary Figure 1 (associated to Figure 1): Validation of the deletion model and lymphatic analysis at 3 months of age**.

A. Quantification of the normalized count for all alpha and beta integrin subunits expressed by the LECs isolated from the meninges (dura), skin (ear) and diaphragm.
B. Schematic of the recombination protocol. $LEC^{WT}$ and $LEC^{\Delta CD49a}$ mice were injected with tamoxifen at 6-7 weeks of age (80mg/kg) for four consecutive days. Efficacy of deletion was measured at 3, 6-7 and 16 months of age.
C. Representative histogram of CD49a expression in dural LECs (left) and BEC (right) in 3 months old $LEC^{WT}$ and $LEC^{\Delta CD49a}$ mice.
D. Quantification of the mean geometric intensity of CD49a expression in LECs (top) and BEC (bottom).
E. Re-analysis of published hippocampal single cell RNA seq data (Sanders et al, Cell 2018) showing mean expression and percentage of cells expressing itga1 across hippocampal cell types.
F. Representative images of the dural lymphatic vessels (Lyve-1; red) along the transverse sinus of young (3 months old) mice.
G. Quantification of the total lymphatic length (i) and lymphatic width (ii) along the transverse sinus of 3 months old male (square) and female (triangle) mice. Mean ± s.e.m. Unpaired t test.
H. Representative images of CSF-injected OVA (green) in the dCLNs of 3 months old mice. Lymph nodes are characterized by the presence of a capsule (Lyve-1; red) and T cells zone (Cd3e; gray).
I. Quantification of the size of the lymph node section. Mean ± s.e.m. Unpaired t test.
J. Quantification of the percentage of Lyve1 coverage. Mean ± s.e.m. Unpaired t test.
K. Quantification of the CD3e coverage. Mean ± s.e.m. Unpaired t test.
L. Quantification of the OVA coverage. Mean ± s.e.m. Unpaired t test.
M. Representative images of CSF injected CFSE labelled naïve CD4 T cells in the T cells zone of the dCLNs of 3 months old mice.
N. Quantification of the density of CFSE+ T cells in the dCLNs. Mean ± s.e.m. Unpaired t test.
O. Representative images of the chemokine Ccl21 (red) in the dural lymphatic vessels (Lyve-1; cyan) of 3 months old mice.
P. Quantification of the Ccl21 coverage of the dural lymphatic vessels. Mean ± s.e.m. Unpaired t test.

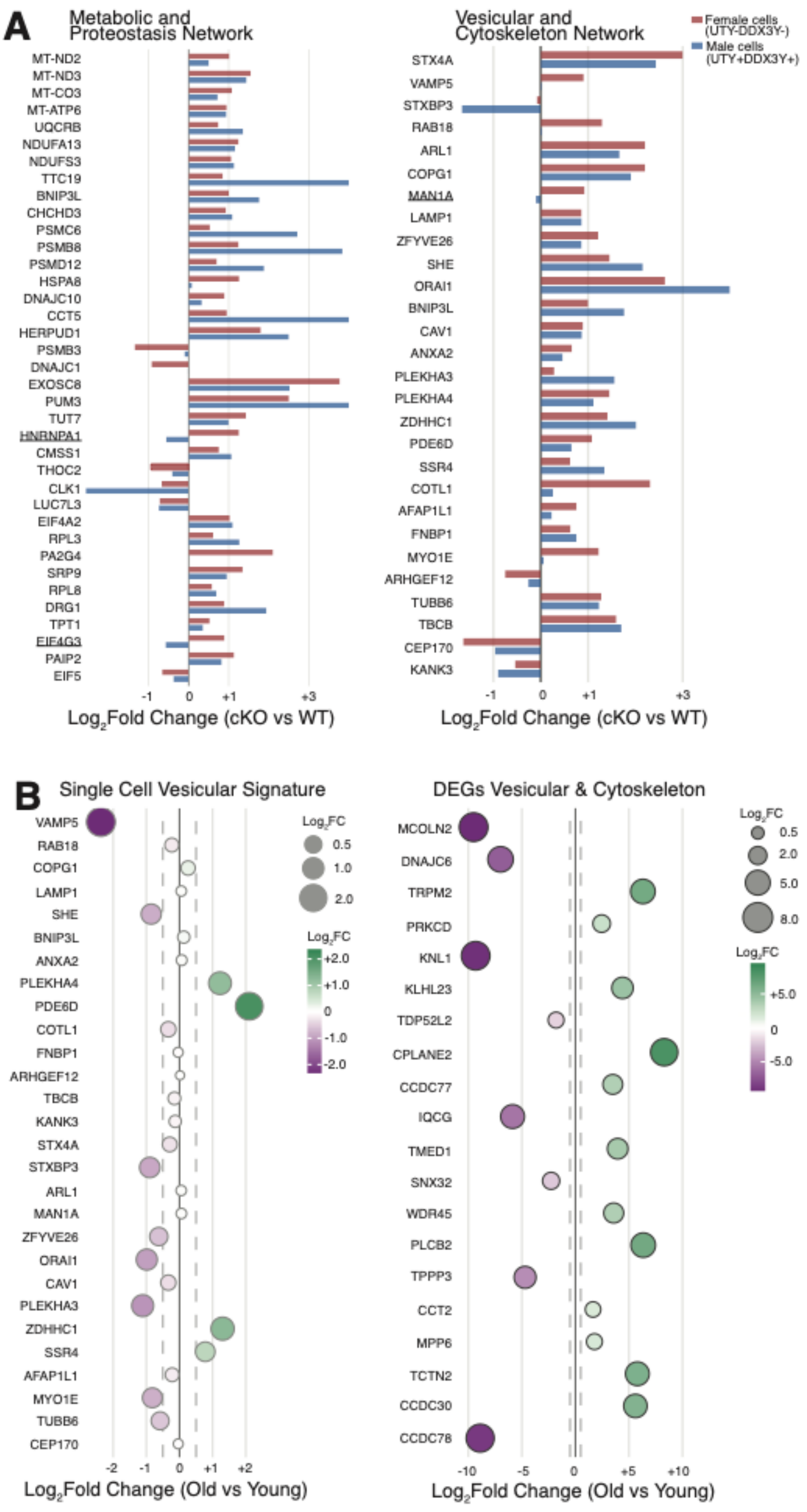
A
Metabolic and Proteostasis Network
MT-ND2
MT-ND3
MT-CO3
MT-ATP6
UQCRB
NDUFA13
NDUFS3
TTC19
BNIP3L
CHCHD3
PSMC6
PSMB8
PSMD12
HSPA8
DNAJC10
CCT5
HERPUD1
PSMB3
DNAJC1
EXOSC8
PUM3
TUT7
HNRNPA1
CMSS1
THOC2
CLK1
LUC7L3
EIF4A2
RPL3
PA2G4
SRP9
RPL8
DRG1
TPT1
EIF4G3
PAIP2
EIF5
-1
0
+1
+3
Log2Fold Change (cKO vs WT)
Vesicular and Cytoskeleton Network
Female cells (UTY-DDX3Y-)
Male cells (UTY+DDX3Y+)
STX4A
VAMP5
STXBP3
RAB18
ARL1
COPG1
MAN1A
LAMP1
ZFYVE26
SHE
ORAI1
BNIP3L
CAV1
ANXA2
PLEKHA3
PLEKHA4
ZDHHC1
PDE6D
SSR4
COTL1
AFAP1L1
FNBP1
MYO1E
ARHGEF12
TUBB6
TBCB
CEP170
KANK3
-1
0
+1
+3
Log2Fold Change (cKO vs WT)
B
Single Cell Vesicular Signature
VAMP5
RAB18
COPG1
LAMP1
SHE
BNIP3L
ANXA2
PLEKHA4
PDE6D
COTL1
FNBP1
ARHGEF12
TBCB
KANK3
STX4A
STXBP3
ARL1
MAN1A
ZFYVE26
ORAI1
CAV1
PLEKHA3
ZDHHC1
SSR4
AFAP1L1
MYO1E
TUBB6
CEP170
-2
-1
0
+1
+2
Log2Fold Change (Old vs Young)
Log2FC
0.5
1.0
2.0
Log2FC
+2.0
+1.0
0
-1.0
-2.0
DEGs Vesicular & Cytoskeleton
MCOLN2
DNAJC6
TRPM2
PRKCD
KNL1
KLHL23
TDP52L2
CPLANE2
CCDC77
IQCG
TMED1
SNX32
WDR45
PLCB2
TPPP3
CCT2
MPP6
TCTN2
CCDC30
CCDC78
-10
-5
0
+5
+10
Log2Fold Change (Old vs Young)
Log2FC
0.5
2.0
5.0
8.0
Log2FC
+5.0
0
-5.0

**Supplementary Figure 2 (associated to Figure 2): Sex specific comparison and cross study analysis of vesicular and cytoskeletal signature**.

A. Bar plots of Log2fold change for individual genes within the Metabolic and Proteostasis network (left) and the Vesicular and Cytoskeleton network (right), shown separately for female cells (*Uty-Ddx3y-*) and male cells (*Uty+Ddx3y+*) illustrating the lack of sex-specific differences in the transcriptional response to CD49a loss.
B. Dot plots of Log2fold change of the genes set related to the Vesicular and Cytoskeleton identified in the single cell RNA seq (left) or DEGs related to Vesicular and Cytokeleton network in the bulk RNA sequencing of aged dural LECs (Da Mesquita et al, 2018).

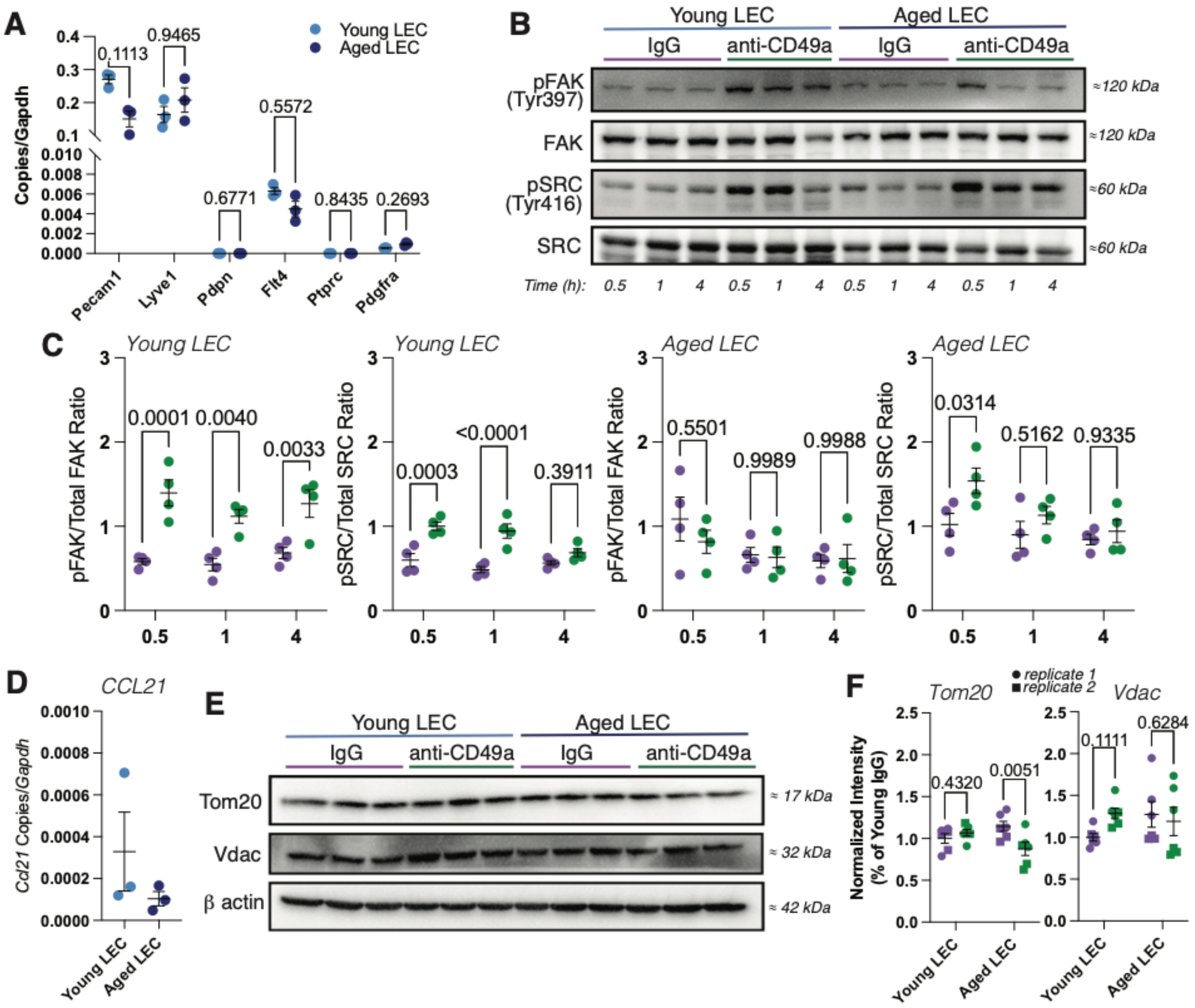
A
Young LEC
Aged LEC
Copies/Gapdh
0.1113
0.9465
0.5572
0.6771
0.8435
0.2693
Pecam1
Lyve1
Pdpn
Flt4
Ptprc
Pdgfra
B
Young LEC
Aged LEC
IgG
anti-CD49a
pFAK (Tyr397)
FAK
pSRC (Tyr416)
SRC
≈120 kDa
≈60 kDa
Time (h):
C
Young LEC
Aged LEC
pFAK/Total FAK Ratio
pSRC/Total SRC Ratio
0.0001
0.0040
0.0033
0.0003
<0.0001
0.3911
0.5501
0.9989
0.9988
0.0314
0.5162
0.9335
D
CCL21
Ccl21 Copies/Gapdh
E
Tom20
Vdac
β actin
≈ 17 kDa
≈ 32 kDa
≈ 42 kDa
F
Tom20
Vdac
replicate 1
replicate 2
Normalized Intensity (% of Young IgG)
0.4320
0.0051
0.1111
0.6284

53

54

**Supplementary Figure 3 (associated to Figure 3): Agonist properties of anti-CD49a antibody and lack of change in mitochondrial load upon chronic CD49a activation**.

A. RT-qPCR quantification of lymphatic/endothelial and lineage marker transcripts, normalized to *Gapdh* in young and aged LECs *in vitro*.
B. Representative western blots of phosphoFAK (Tyr397), total FAK, phosphoSRC (Tyr416), total SRC in young and aged LEC treated with IgG or anti-CD49a.
C. Quantification of pFAK/total FAK and pSRC/total SRC ratio over time following IgG or anti-CD49a treatment in young (left two plots) and aged (right two plots) LEC. Each dot represents a biological replicate of the cell treatment. Mean $\pm$ s.e.m. Two-way ANOVA with Sidak's multiple comparisons test.
D. RT-qPCR of *Ccl21* transcript, normalized to *Gapdh*, in young and aged LECs *in vitro*. Mean $\pm$ s.e.m.
E. Representative western blots of Tom20, VDAC (mitochondrial mass markers) and beta actin in young and aged LECS treated with IgG and anti-CD49a.
F. Quantification of the normalized expression of Tom20 and Vdac expressed as a percentage of the young IgG condition. Mean $\pm$ s.e.m. Two-way ANOVA with Uncorrected Fisher's LSD

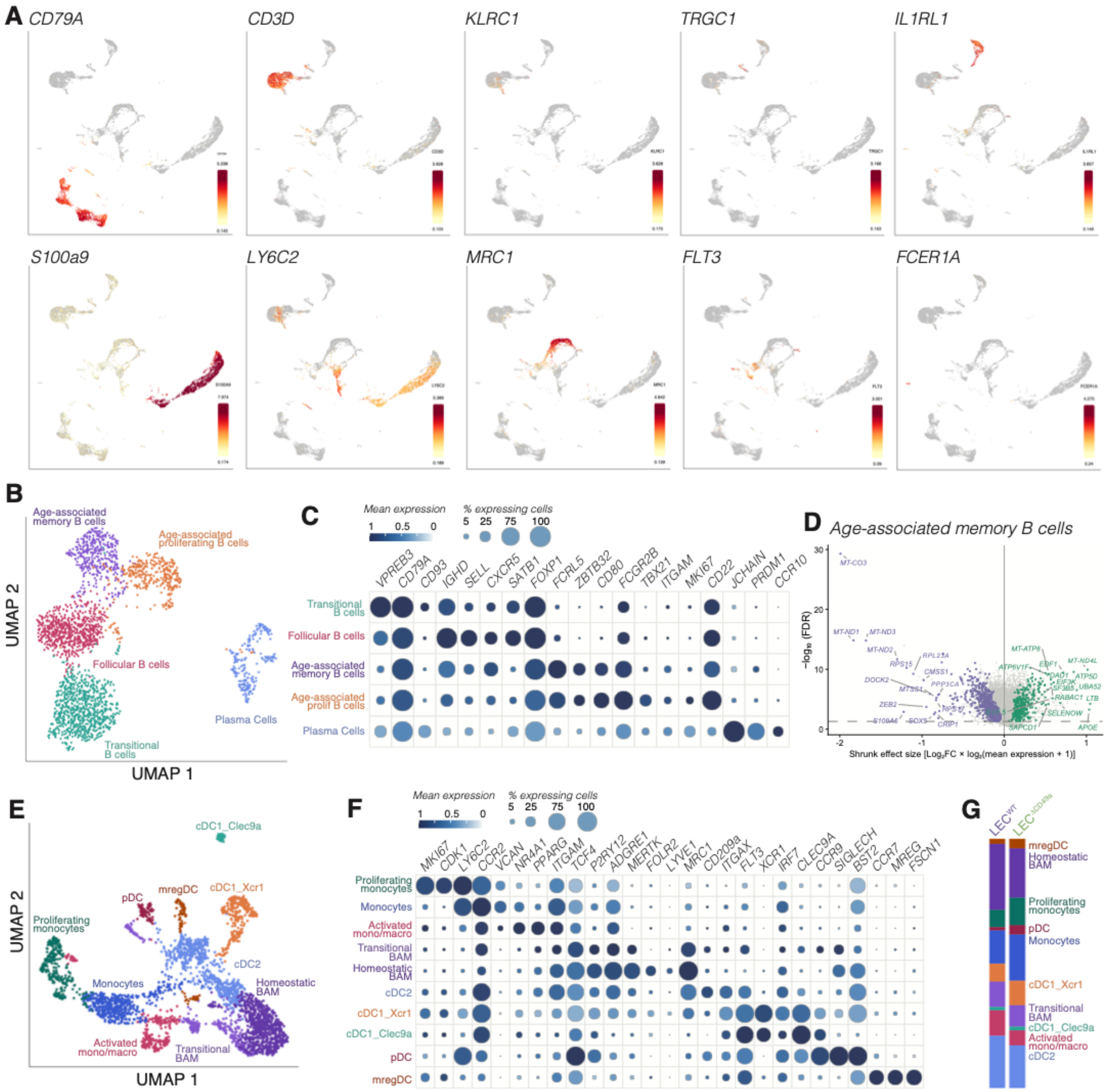
A
CD79A
CD3D
KLRC1
TRGC1
IL1RL1
S100a9
LY6C2
MRC1
FLT3
FCER1A
B
Age-associated memory B cells
Age-associated proliferating B cells
Follicular B cells
Transitional B cells
Plasma Cells
UMAP 2
UMAP 1
C
Mean expression
1 0.5 0
% expressing cells
5 25 75 100
VPREB3
CD79A
CD93
IGHD
SELL
CXCR5
SATB1
FOXP1
FCRL5
ZBTB32
CD80
FCGR2B
TBX21
ITGAM
MKI67
CD22
JCHAIN
PRDM1
CCR10
Transitional B cells
Follicular B cells
Age-associated memory B cells
Age-associated prolif B cells
Plasma Cells
D
Age-associated memory B cells
−log10 (FDR)
Shrunk effect size [Log2FC × log2(mean expression + 1)]
MT-CO3
MT-ND1
MT-ND3
MT-ND2
RPS15
RPL23A
CMSS1
PPP3CA
DOCK2
MTSS1
ZEB2
S100A6
SOX5
CRIP1
MT-ATP8
ATP6V1F
EDF1
MT-ND4L
ATP5D
UBA52
RABAC1
LTB
SELENOW
SAPCD1
APOE
E
cDC1_Clec9a
pDC
mregDC
cDC1_Xcr1
Proliferating monocytes
cDC2
Monocytes
Homeostatic BAM
Activated mono/macro
Transitional BAM
UMAP 2
UMAP 1
F
Mean expression
1 0.5 0
% expressing cells
5 25 75 100
MKI67
CDK1
LY6C2
CCR2
VCAN
NR4A1
PPARG
ITGAM
TCF4
P2RY12
ADGRE1
MERTK
FOLR2
LYVE1
MRC1
CD209a
ITGAX
FLT3
XCR1
IRF7
CLEC9A
CCR9
SIGLECH
BST2
CCR7
MREG
FSCN1
Proliferating monocytes
Monocytes
Activated mono/macro
Transitional BAM
Homeostatic BAM
cDC2
cDC1_Xcr1
cDC1_Clec9a
pDC
mregDC
G
LEC^WT
mregDC
Homeostatic BAM
Proliferating monocytes
pDC
Monocytes
cDC1_Xcr1
Transitional BAM
cDC1_Clec9a
Activated mono/macro
cDC2

**Supplementary Figure 4 (associated to Figure 4): Transcriptomic changes in dural immune cells upon CD49a loss in aged LECs**.

A. UMAP feature plots showing normalized expression of canonical lineage markers genes used to annotate major immune cell populations in the aged dura. Color scale indicated normalized expression level.
B. UMAP of the B cell subset, colored by annotated subset type.
C. Dot plot of canonical marker genes defining each B cell subpopulation from (B). Dot color indicates mean expression while dot size indicated percentage of expressing cells.
D. Volcano plot of differentially expressed genes in the Age-associated memory B cell cluster, plotting shrunken effect size against -log10(FDR). Genes upregulated in $LEC^{\Delta CD49a}$ are shown in green; genes downregulated are shown in purple.
E. UMAP of the myeloid subset, colored by annotated subtype.
F. Dot plot of canonical marker genes defining each myeloid subpopulation from (E), formatted as in (C).
G. Stacked bar plot of the relative proportion of each myeloid subpopulation in $LEC^{WT}$ versus $LEC^{\Delta CD49a}$.

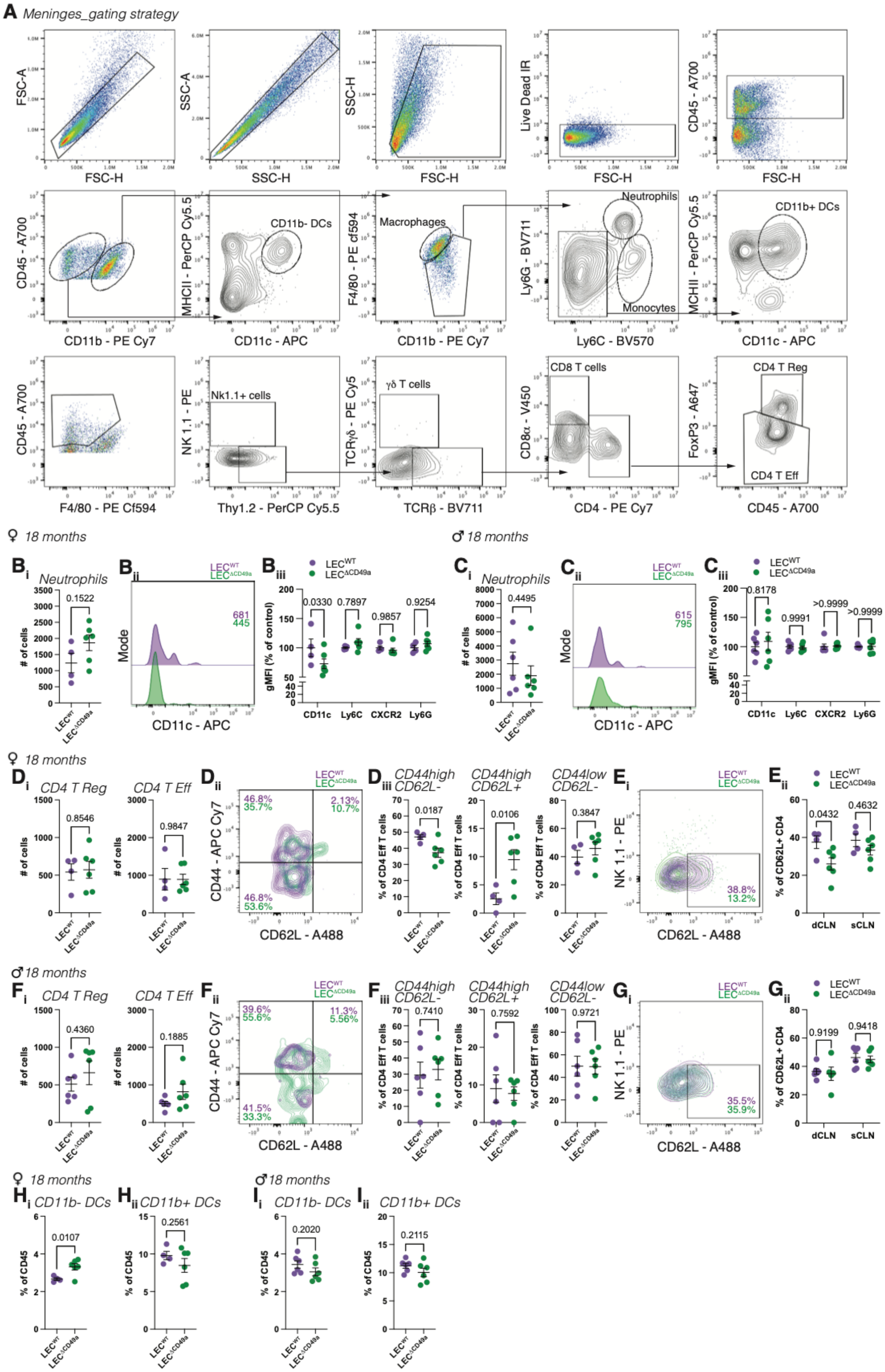

A Meninges_gating strategy
FSC-A
FSC-H
SSC-A
SSC-H
Live Dead IR
CD45 - A700
CD11b - PE Cy7
MHCII - PerCP Cy5.5
CD11b- DCs
CD11c - APC
F4/80 - PE cf594
Macrophages
Ly6G - BV711
Neutrophils
Monocytes
Ly6C - BV570
CD11b+ DCs
F4/80 - PE Cf594
NK 1.1 - PE
Nk1.1+ cells
Thy1.2 - PerCP Cy5.5
TCRγδ - PE Cy5
γδ T cells
TCRβ - BV711
CD8α - V450
CD8 T cells
CD4 - PE Cy7
FoxP3 - A647
CD4 T Reg
CD4 T Eff
♀ 18 months
♂ 18 months
Bi Neutrophils
### of cells
0.1522
Bii
Mode
681
445
Biii
LEC^WT
LEC^ΔCD49a
gMFI (% of control)
0.0330
0.7897
0.9857
0.9254
CD11c
Ly6C
CXCR2
Ly6G
Ci Neutrophils
0.4495
Cii
615
795
Ciii
0.8178
0.9991
>0.9999
>0.9999
Di CD4 T Reg
0.8546
CD4 T Eff
0.9847
Dii
CD44 - APC Cy7
CD62L - A488
46.8%
35.7%
2.13%
10.7%
46.8%
53.6%
Diii CD44high CD62L-
% of CD4 Eff T cells
0.0187
CD44high CD62L+
0.0106
CD44low CD62L-
0.3847
Ei
38.8%
13.2%
Eii
% of CD62L+ CD4
0.0432
0.4632
dCLN
sCLN
Fi CD4 T Reg
0.4360
CD4 T Eff
0.1885
Fii
39.6%
55.6%
11.3%
5.56%
41.5%
33.3%
Fiii CD44high CD62L-
0.7410
CD44high CD62L+
0.7592
CD44low CD62L-
0.9721
Gi
35.5%
35.9%
Gii
0.9199
0.9418
Hi CD11b- DCs
% of CD45
0.0107
Hii CD11b+ DCs
0.2561
Ii CD11b- DCs
0.2020
Iii CD11b+ DCs
0.2115

**Supplementary Figure 5 (associated to Figure 4): Flow cytometry analysis of the aged dural immune cells in** LEC$^{\Delta CD49a}$ **mice**.

A. Dural immune cell gating strategy.
B. Quantification of the number of dural neutrophils (i), representative histogram of CD11c expression in dural neutrophils (ii) and quantification of the geometric mean fluorescence intensity of Cd11c, Ly6c, Ly6c and Cxcr2 expression by dural neutrophils (iii) in aged female LEC$^{\Delta CD49a}$ mice compared to LEC$^{WT}$. Mean $\pm$ s.e.m. Welch's t test (i); two-way ANOVA with Sidak's multiple comparisons test (iii).
C. Quantification of the number of dural neutrophils (i), representative histogram of Cd11c expression in dural neutrophils (ii) and quantification of the geometric mean fluorescence intensity of Cd11c, Ly6c, Ly6g and Cxcr2 expression by dural neutrophils (iii) in aged male LEC$^{\Delta CD49a}$ mice compared to LEC$^{WT}$. Mean $\pm$ s.e.m. Welch's t test (i); two-way ANOVA with Sidak's multiple comparisons test (iii).
D. Quantification of the number of regulatory (left) or effector (right) CD4 T cells (i), representative contour plot of Cd62l and Cd44 expression by dural effector CD4 T cells (ii), and quantification of the percentage of CD4 T cells of the indicated phenotype (iii) between aged LEC$^{\Delta CD49a}$ and LEC$^{WT}$ female mice. Mean $\pm$ s.e.m. Welch's t test.
E. Representative contour plot of Cd62l expression by CD4 T cells (i) and quantification of the percentage of Cd62l+ CD4 T cells in the dCLNs and sCLNs of LEC$^{\Delta CD49a}$ aged female mice compared to LEC$^{WT}$. Mean $\pm$ s.e.m. Two-way ANOVA with Sidak's multiple comparisons test.
F. Quantification of the number of regulatory (left) or effector (right) CD4 T cells (i), representative contour plot of Cd62l and Cd44 expression by dural effector CD4 T cells (ii), and quantification of the percentage of CD4 T cells of the indicated phenotype (iii) between aged LEC$^{\Delta CD49a}$ and LEC$^{WT}$ male mice. Mean $\pm$ s.e.m. Welch's t test.
G. Representative contour plot of Cd62l expression by CD4 T cells (i) and quantification of the percentage of Cd62l+ CD4 T cells in the dCLNs and sCLNs of LEC$^{\Delta CD49a}$ aged male mice compared to LEC$^{WT}$. Mean $\pm$ s.e.m. Two-way ANOVA with Sidak's multiple comparisons test.
H. Quantification of the percentage of Cd11b-DCs (i) and Cd11b+DCs (ii) amongst Cd45+ cells in aged female LEC$^{\Delta CD49a}$ versus LEC$^{WT}$. Mean $\pm$ s.e.m. Welch's t test.
I. Quantification of the percentage of Cd11b-DCs (i) and Cd11b+DCs (ii) amongst Cd45+ cells in aged male LEC$^{\Delta CD49a}$ versus LEC$^{WT}$. Mean $\pm$ s.e.m. Welch's t test.

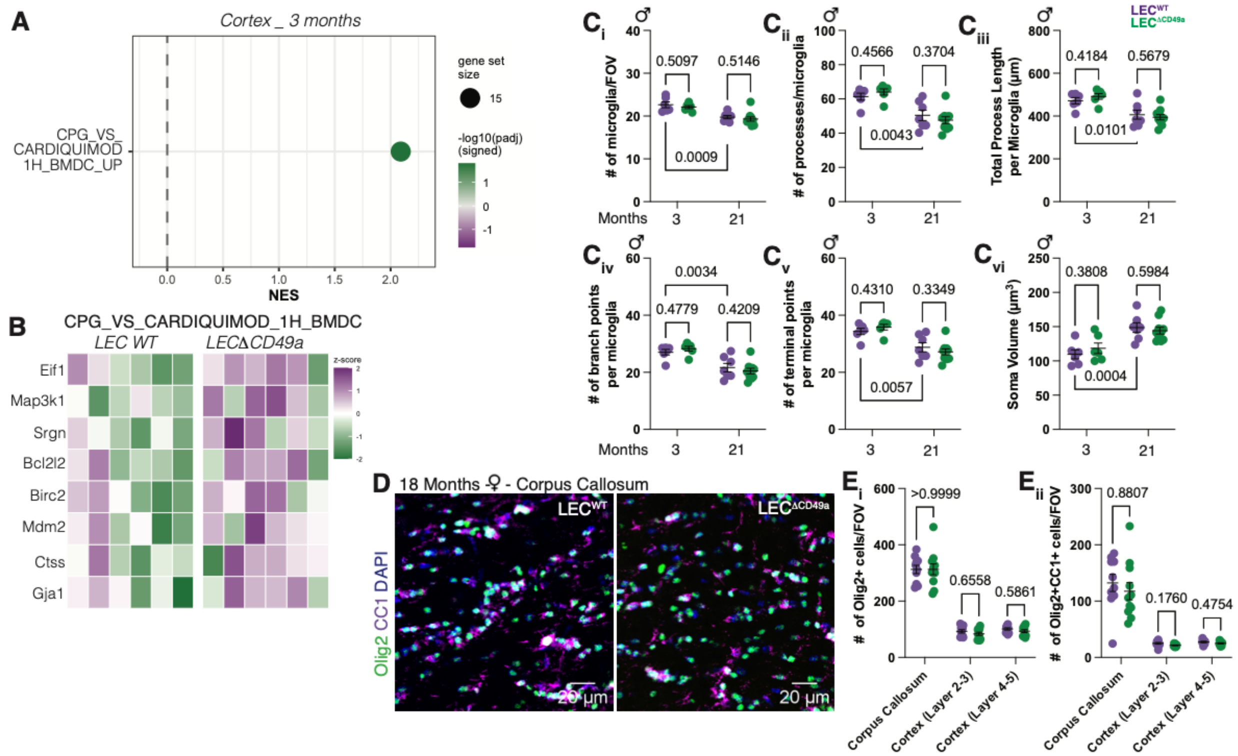

A
Cortex _ 3 months
CPG_VS_ CARDIQUIMOD 1H_BMDC_UP
gene set size
15
-log10(padj) (signed)
NES
B
CPG_VS_CARDIQUIMOD_1H_BMDC
LEC WT
LECΔCD49a
Eif1
Map3k1
Srgn
Bcl2l2
Birc2
Mdm2
Ctss
Gja1
z-score
C i
### of microglia/FOV
0.5097
0.5146
0.0009
Months
C ii
### of processes/microglia
0.4566
0.3704
0.0043
C iii
Total Process Length per Microglia (µm)
0.4184
0.5679
0.0101
LEC^WT
LEC^ΔCD49a
C iv
### of branch points per microglia
0.0034
0.4779
0.4209
C v
### of terminal points per microglia
0.4310
0.3349
0.0057
C vi
Soma Volume (µm³)
0.3808
0.5984
0.0004
D 18 Months ♀ - Corpus Callosum
LEC^WT
LEC^ΔCD49a
Olig2 CC1 DAPI
20 µm
E i
### of Olig2+ cells/FOV
>0.9999
0.6558
0.5861
E ii
### of Olig2+CC1+ cells/FOV
0.8807
0.1760
0.4754
Corpus Callosum
Cortex (Layer 2-3)
Cortex (Layer 4-5)


130

131

**Supplementary Figure 6 (associated to Figure 5): Gene set enrichment in young mice, male microglial morphology, and oligodendrocyte lineage quantifications**.

A. Gene set enrichment analysis (GSEA) of Nanostring Neuroinflammation panel data from cortex of 3 months old mice comparing $LEC^{\Delta CD49a}$ to $LEC^{WT}$.
B. Heatmap of z-scored expression for individual genes within the gene set identified in (A), comparing $LEC^{\Delta CD49a}$ to $LEC^{WT}$. Each column represents an individual biological replicate, color scale indicated row-wise z-score.
C. Quantification of the number of microglia (i), number of processes per microglia (ii), total length of processes per microglial (iii), number of branch points per microglia (iv), number of terminal points per microglia (v) and microglial soma volume (vi) in young and aged $LEC^{WT}$ and $LEC^{\Delta CD49a}$ male mice. Mean $\pm$ s.e.m. Two-way ANOVA with Uncorrected Fisher's LSD.
D. Representative images of the corpus callosum from female $LEC^{WT}$ and $LEC^{\Delta CD49a}$ at 18 months, stained for Olig2 (green), Cc1 (magenta) and Dapi (blue).
E. Quantification of the density of Olig2+ cell (i) and Olig2+Cc1+ (mature oligodendrocyte) per field of view in corpus callosum and cortical layers comparing $LEC^{WT}$ and $LEC^{\Delta CD49a}$ aged female mice. Mean $\pm$ s.e.m. Two-way ANOVA with Sidak's multiple comparison test.

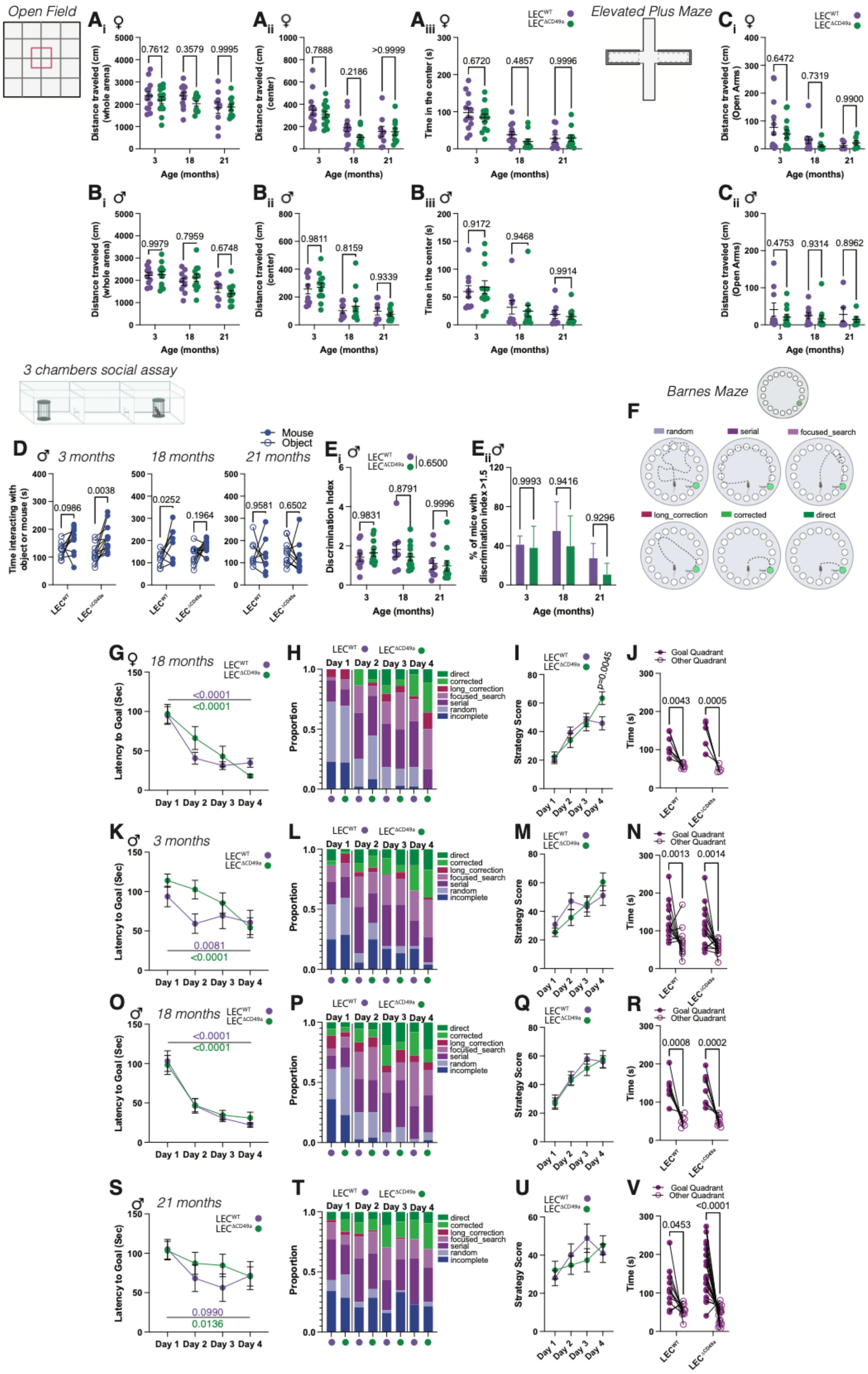
Open Field
Elevated Plus Maze
3 chambers social assay
Barnes Maze
3 months
18 months
21 months
Mouse
Object
random
serial
focused_search
long_correction
corrected
direct
incomplete
Goal Quadrant
Other Quadrant
Age (months)
Distance traveled (cm) (whole arena)
Distance traveled (cm) (center)
Time in the center (s)
Distance traveled (cm) (Open Arms)
Time interacting with object or mouse (s)
Discrimination Index
% of mice with discrimination index >1.5
Latency to Goal (Sec)
Proportion
Strategy Score
Time (s)
Day 1
Day 2
Day 3
Day 4
p=0.0045

**Supplementary Figure 7 (associated to Figure 6) : Open field and elevated plus maze analysis, male behavior data and female Barnes maze at 18 months of age**.

A. Quantification of the total distance (i), distance travelled in the center (ii) and time spent in the center of the open field for female $LEC^{\Delta CD49a}$ compared to $LEC^{WT}$ at 3, 18 and 21 months of age. Mean ± s.e.m. Two-way ANOVA with Sidak's multiple comparison test.
B. Quantification of the total distance (i), distance travelled in the center (ii) and time spent in the center of the open field for male $LEC^{\Delta CD49a}$ compared to $LEC^{WT}$ at 3, 18 and 21 months of age. Mean ± s.e.m. Two-way ANOVA with Sidak's multiple comparison test.
C. Quantification of the total distanced travelled in the open arms for female (top) and male (bottom) $LEC^{\Delta CD49a}$ compared to $LEC^{WT}$ at 3, 18 and 21 months of age. Mean ± s.e.m. Two-way ANOVA with Sidak's multiple comparison test.
D. Quantification of the time spent interacting with a mouse or inanimate object in the 3-chamber social assay at 3, 18 and 21 months of age comparing $LEC^{WT}$ and $LEC^{\Delta CD49a}$ male mice. Mean ± s.e.m. Two-way ANOVA with Sidak's multiple comparisons test.
E. (i) Quantification of the sociability ratio between $LEC^{WT}$ and $LEC^{\Delta CD49a}$ male mice at 3, 18 and 21 months of age. Mean ± s.e.m. Two-way ANOVA with Sidak's multiple comparisons test. (ii) Quantification of the percentage of mice with a discrimination index superior to 1.5 in $LEC^{WT}$ and $LEC^{\Delta CD49a}$ male mice at 3, 18 and 21 months of age. Mean ± s.e.m. Two-way ANOVA with Sidak's multiple comparisons test.
F. Schematic of the strategy categories used for the quantification.
G. Quantification of the latency to reach the goal hole during the training portion of the Barnes maze in 18 months old females. N=5/8 mice/group. Mean ± s.e.m. Two-way ANOVA with Sidak's multiple comparisons test.
H. Quantification of the proportion of strategy usage in 18 months old females.
I. Quantification of the strategy score over the training days in 18 months old females. N=5/8 mice/group. Mean ± s.e.m. Two-way ANOVA with Sidak's multiple comparisons test.
J. Quantification of the time spent in the goal quadrant compared to the other quadrants during the test phase of the Barnes maze in 18 months old females. N=5/8 mice/group. Mean ± s.e.m. Two-way ANOVA with Sidak's multiple comparisons test.
K. Quantification of the latency to reach the goal hole during the training portion of the Barnes maze in 3 months old males. N=13/15 mice/group. Mean ± s.e.m. Two-way ANOVA with Sidak's multiple comparisons test.

L. Quantification of the proportion of strategy usage in 3 months old males.

M. Quantification of the strategy score over the training days in 3 months old male. N=13/15 mice/group. Mean ± s.e.m. Two-way ANOVA with Sidak's multiple comparisons test.

N. Quantification of the time spent in the goal quadrant compared to the other quadrants during the test phase of the Barnes maze in 3 months old male. N=13/15 mice/group. Mean ± s.e.m. Two-way ANOVA with Sidak's multiple comparisons test.

O. Quantification of the latency to reach the goal hole during the training portion of the Barnes maze in 18 months old males. N=9/12 mice/group. Mean ± s.e.m. Two-way ANOVA with Sidak's multiple comparisons test.

P. Quantification of the proportion of strategy usage in 18 months old males.

Q. Quantification of the strategy score over the training days in 18 months old males. N=9/12 mice/group. Mean ± s.e.m. Two-way ANOVA with Sidak's multiple comparisons test.

R. Quantification of the time spent in the goal quadrant compared to the other quadrant during the test phase of the Barnes maze in 18 months old males. N=9/12 mice/group. Mean ± s.e.m. Two-way ANOVA with Sidak's multiple comparisons test.

S. Quantification of the latency to reach the goal hole during the training portion of the Barnes maze 21 months old males. N=11/21 mice/group. Mean ± s.e.m. Two-way ANOVA with Sidak's multiple comparisons test.

T. Quantification of the proportion of strategy usage in 21 months old males.

U. Quantification of the strategy score over the training days in 21 months old males. N=11/21 mice/group. Mean ± s.e.m. Two-way ANOVA with Sidak's multiple comparisons test.

V. Quantification of the time spent in the goal quadrant compared to the other quadrants during the test phase of the Barnes maze in 21 months old males. N=11/21 mice/group. Mean ± s.e.m. Two-way ANOVA with Sidak's multiple comparisons test.
